# Enrichment analysis of GWAS data in autoimmunity delineates the multiple sclerosis-Epstein Barr virus association

**DOI:** 10.1101/2021.06.06.447253

**Authors:** Rosella Mechelli, Renato Umeton, Virginia Rinaldi, Gianmarco Bellucci, Rachele Bigi, Daniela F. Angelini, Gisella Guerrera, Sundararajan Srinivasan, Silvia Romano, Maria Chiara Buscarinu, Raffaella Pizzolato Umeton, Eleni Anastasiadou, Pankaj Trivedi, Arianna Fornasiero, Michela Ferraldeschi, IMSGC and WTCCC2, Diego Centonze, Antonio Uccelli, Dario Di Silvestre, Pier Luigi Mauri, Paola de Candia, Giuseppe Matarese, Sandra D’Alfonso, Luca Battistini, Cinthia Farina, Roberta Magliozzi, Richard Reynolds, Marco Salvetti, Giovanni Ristori

**Author notes:** **Correspondence: Rosella Mechelli**, San Raffaele Roma Open University and IRCCS San Raffaele Pisana, Rome, Italy, Via di Val Cannuta, 247, 00166 Rome (Italy), **Giovanni Ristori**, Centre for Experimental Neurological Therapies (CENTERS), Department of Neurosciences, Mental Health and Sensory Organs, Sapienza University, Rome, Via di Grottarossa, 1035-1039, 00189 Rome (Italy), **Marco Salvetti**, Centre for Experimental Neurological Therapies (CENTERS), Department of Neurosciences, Mental Health and Sensory Organs, Sapienza University, Rome, Via di Grottarossa, 1035-1039, 00189 Rome (Italy). These authors contributed equally to this work as co-first authors (Rosella Mechelli, Renato Umeton).

## Abstract

We exploited genetic information to assess non-genetic influences in autoimmunity. We isolated gene modules whose products physically interact with environmental exposures related to autoimmunity, and analyzed their nominal statistical evidence of association with autoimmune and non-autoimmune diseases in genome-wide association studies (GWAS) data. Epstein Barr virus (EBV) and other Herpesviruses interactomes emerged as specifically associated with multiple sclerosis (MS), possibly under common regulatory mechanisms. Analyses of MS blood and brain transcriptomes, cytofluorimetric studies of endogenous EBV-infected lymphoblastoid lines, and lesion immunohistochemistry, confirmed a dysregulation of MS-associated EBV interactors, suggesting their contribution to CD40 signaling alterations in MS. These interactors resulted enriched in modules from inherited axonopathies-causing genes, supporting a link between EBV and neurodegeneration in MS, in accord with the observed transcriptomic dysregulations in MS brains. They were also enriched with top-ranked pharmaceutical targets prioritized on a genetic basis. This study delineates a disease-specific influence of herpesviruses on MS biology.

## INTRODUCTION

By definition, multifactorial diseases are thought to be caused by heritable and nonheritable factors. Genome-wide association studies (GWAS) provide information about the former and sero-epidemiological studies do the same for the latter. Both types of studies are subject to interpretation difficulties. One limitation of GWAS derives from the fragmentation of the information they provide, i.e. lists of single variables, mostly of small effect size, whose putative functions are difficult to reunite and reorganize in one or few disease processes (Boyle et al. 2017; Califano et al. 2012; Li et al. 2019). For seroepidemiological studies, the limitation comes from the correlative and associative nature of most of the investigations on environmental exposures (Fischbach 2018). However, GWAS can inform about the causal relevance of associations between environmental exposures and disease. Reciprocally, extracting value from the study of the interaction between relevant exposures and predisposing genes may contribute to the interpretation of GWAS, in a virtuous ‘genes-to-environment-to-genes-again’ iterative learning process (Tam et al. 2019; Visscher et al. 2017).

Previous work has shown the potential relevance of GWAS data analyses that focus on the combined effects of many loci of small effect-size (Beecham et al. 2013; Purcell et al. 2009; Saez-Atienzar et al. 2021). Among different efforts to study biologically meaningful combinations of genes, we devised a “candidate interactome” approach to investigate which gene-environment interactions may associate with the development of multiple sclerosis (MS), a multifactorial disease of the central nervous system (CNS) that affects young adults, resulting in significant disability and socioeconomic burden for both affected persons and society. In GWAS data, we looked for statistical enrichments of associations among modules of genes whose products interact with environmental factors of plausible, uncertain, or unlikely relevance for MS pathogenesis (“candidate interactomes”) (Mechelli et al. 2013). This approach can be considered akin to Mendelian randomization, which exploits genetic data to improve causal inference in epidemiology (Davey Smith and Hemani 2014). The results provided a preliminary suggestion about the possible relevance of the interaction between host genotype and Epstein Barr virus (EBV) in MS. However, this study did not investigate the disease-specificity and the biological implications of the observed associations.

Integrating GWAS information with transcriptional regulatory data and protein interactions provides plausible models of disease pathogenesis (Baird et al. 2021; Consortium 2019a). Elegant studies are resolving the functional implications of individual disease-associated variants (Gregory et al. 2012; Steri et al. 2017). However, given the complex nature of multifactorial diseases (Wray et al. 2018), and particularly for translation into clinical benefit (Zeggini et al. 2019), a general understanding of how gene-environment interactions generate gene expression alterations is also necessary (Gallagher and Chen-Plotkin 2018).

Finally, it is known that drug targets having genetic associations with a disease significantly increase the probability of success in drug development (Nelson et al. 2015). In our approach, since the genetic associations reflect environmental exposures, the therapeutic relevance may be increased by the opportunity of targeting not only the genetic component but also the exposure. Moreover, in many autoimmune diseases, where the therapeutic scenario is rapidly changing, it is important to understand which targets may be of etiologic relevance in order to design an appropriate therapeutic sequencing.

Taking advantage of the increasing coverage and accuracy of protein-protein interaction data (Menche et al. 2015), and of the availability of additional MS-GWAS data, in this study we replicated and refined the previous findings (Mechelli et al. 2013), extending the “candidate interactome “analysis to GWAS in other multifactorial conditions. This also allowed the assessment of the disease-specificity of the results. We also investigated the functional implications of the observed associations and the distribution of the therapeutic validity (Fang et al. 2019) of the genes highlighted by the “candidate interactome” approach. Overall, this study outlined an influence of EBV and other Herpesviruses on the biology of MS, which differentiates this disease from other immune-mediated disorders. This influence involves immunological aspects but may also relate to the neurodegenerative component of the disease.

## RESULTS

### Selection and construction of interactome modules

We isolated modules of genes whose products are known to physically interact (“interactomes”) with environmental exposures or in biological processes of plausible, uncertain, or unlikely relevance for the following autoimmune and non-autoimmune complex disorders: MS, type 1 and type 2 diabetes, rheumatoid arthritis, Crohn disease, celiac disease, bipolar disorder, hypertension, coronary artery disease. We considered 20 interactomes (Figure 1A) (10 related to environmental exposures and 10 to biological processes): interactomes of 9 viruses [Epstein-Barr virus (EBV), Hepatitis B virus (HBV), Cytomegalovirus (CMV), Human Herpes virus 8 (HHV8), JC virus (JCV), Human immunodeficiency virus (HIV), Hepatitis C virus (HCV), H1N1-influenza (H1N1), Polyomavirus], 1 bacterium (Chlamydia), 9 protein/process hubs [Autoimmune regulator (AIRE), Vitamin D receptor (VDR), Aryl hydrocarbon receptor (AHR), proteins targeted by 70 innate immune-modulating viral open reading frames from 30 viral species (VIRORF), human innate immunity interactome for type I interferon (h-IFN), histone deacetylases (HDAC), Sirtuin 1 (SIRT1), Sirtuin 7 (SIRT7), Inflammasome] and 1 Human-microRNA targets (human miRNA-mRNA) network. The manually curated interactomes (CMV, EBV, HBV, HHV8 and JCV) and those obtained from publicly available databases (AHR, AIRE and VDR), that we had already selected in our previous work (Mechelli et al. 2013), were updated. We constructed the Inflammasome interactome by analyzing the literature and selecting all the physical-direct interactions among inflammasome proteins and other cellular proteins. The HDACs interactome was obtained from a unique source where different methods had been used to highlight protein-protein interactions among the 11 known HDAC enzymes and other cellular proteins (Joshi et al. 2013). We considered also other deacetylase enzymes belonging to the sirtuin family (SIRT1 and SIRT7), whose interactomes were obtained from BioGRID database (Stark et al. 2006). For the human miRNA-mRNA interactome we used miRecords, taking into consideration only experimentally verified miRNA human targets (Xiao et al. 2009). The Chlamydia interactome was obtained from Mirrashidi et al., that used affinity purification and mass spectrometry approaches to highlight interactions among bacterial and cellular proteins (Mirrashidi et al. 2015). Finally, we used VirusMentha, a protein-protein interaction database specific for virus-host interactions (Calderone et al. 2015), to obtain Polyomavirus interactome.

**Figure 1:**
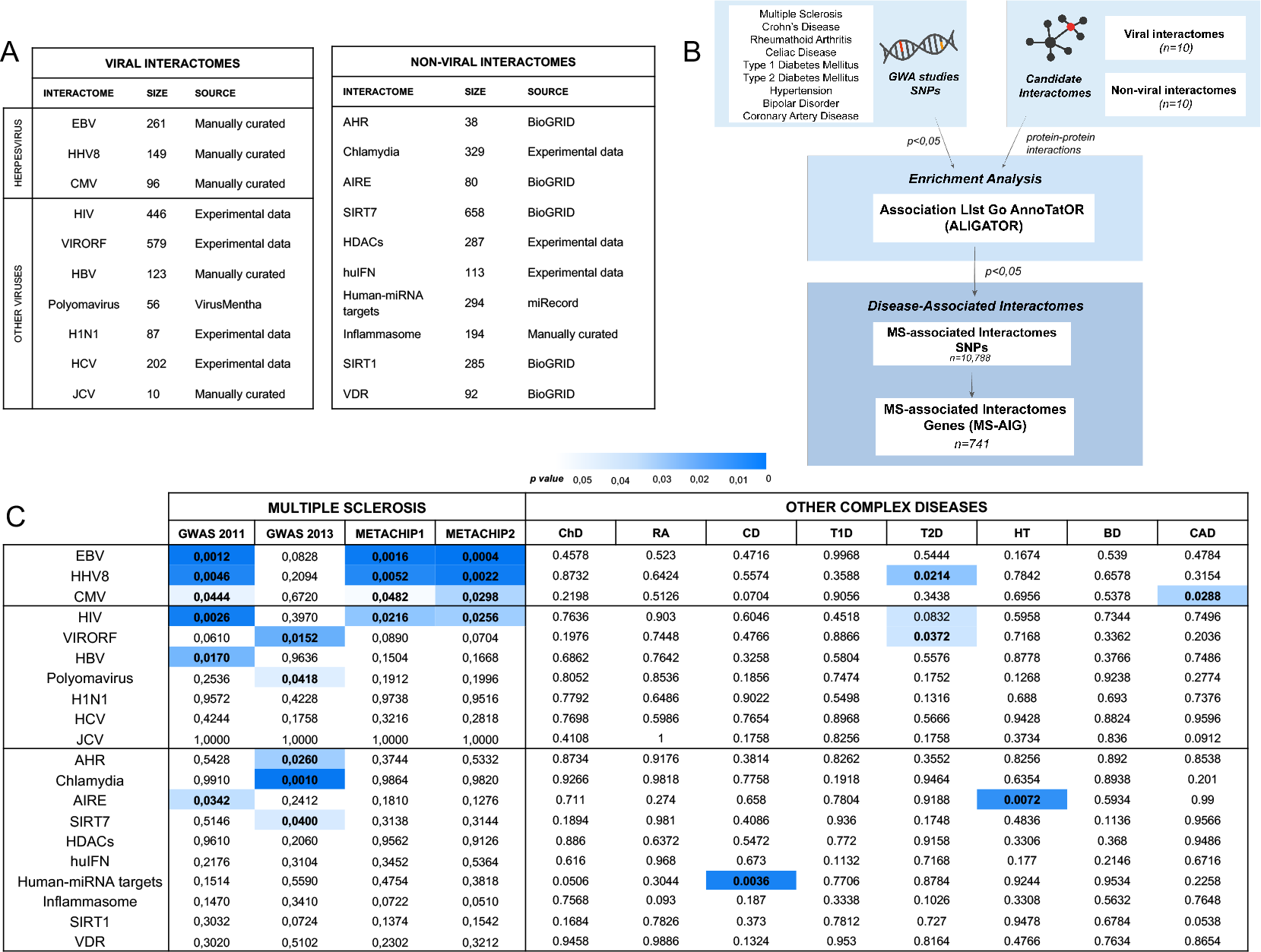
Disease-associated interactomes. (A) Schematic representation of candidate-interactome analysis. (B) List of interactomes and related sizes and sources. (C) Heatmap represents the interactomes and related strength of associations with diseases. The statistically relevant associations are indicated in blue, with a color gradient from white (p> 0.05) to blue (p< 0.01). ChD= Crohn disease; RA= rheumatoid arthritis; CD= celiac disease; T1D= type 1 diabetes; T2D= type 2 diabetes; HT= hypertension; BD= bipolar disorder; CAD= coronary artery disease; AHR= Aryl hydrocarbon receptor; AIRE= autoimmune regulator; BioGRID= Biological General Repository for Interaction Datasets; CMV= Cytomegalovirus; EBV= Epstein-Barr virus; HBV= Hepatitis B virus; HCV= Hepatitis C virus; HDACs=Histone deacetylases; HHV8= Human Herpesvirus 8; HIV= Human Immunodeficiency virus; H1N1= Influenza A virus; hu-IFN= human innate immunity interactome for type I interferon; Human-miRNA targets= gene targets for human miRNA; Inflammasome= multiprotien complex responsible for activation of inflammatory processes and pyroptosis; JCV= JC virus; SIRT1= Sirtuin 1; SIRT7= Sirtuin 7; VIRORF= Virus Open Reading Frame; VDR= vitamin D receptor;

The MS-GWAS data came from the data set published in 2011 (Sawcer et al. 2011) and from Immunochip data (Beecham et al. 2013). To globally evaluate the contribution of all MS-associated SNPs, we constructed two combinations of the MS-GWAS 2011 and MS-GWAS 2013: in the first combination, METACHIP1, MS-GWAS 2011 data were considered in case of overlap (i.e., if both chips had a SNP in the same position, the MS-GWAS 2011 p-value was preferred); in the second combination, METACHIP2, MS-GWAS 2013 data were preferred in case of overlap (Figure S1). The GWAS data in the other diseases were from the Wellcome Trust Case Control Consortium 2 (Consortium 2007).

From GWAS datasets, we selected significant SNPs associated with MS and with the other complex diseases, considering a p-value cut-off of association with each disease at 0.05. We investigated their statistical enrichment of association within each one of the 20 selected interactomes using Association LIst Go AnnoTatOR (ALIGATOR; see workflow in Figure 1B). Spearman’s coefficients were calculated to evaluate linear correlation between the size of single interactomes and their cumulative p-value of association with the diseases, but no correlation was found in any case, indicating that the size of interactomes does not influence discovery (Figure S2-3).

### Herpesvirus interacting proteins are enriched in MS risk genes

When analyzing the frequency of disease-associated genes from distinct GWAS studies in the 20 interactomes, viral interactomes were consistently associated with MS (Figure 1 C). The three herpes viruses studied showed statistical significance with good consistency throughout the analyses (levels of significance: EBV>HHV8>CMV), except for MS GWAS 2013. Given the design of GWAS 2013 (SNPs selected from immune-related loci shared by autoimmune disorders), our result may suggest that herpesviruses do not associate to MS through genetic traits common to other autoimmune diseases. In fact, the herpes virus association characterized MS with respect to all other conditions, including immune-mediated disorders like rheumatoid arthritis, Crohn disease and celiac disease (Figure 1C). Significant associations were sparse in non-MS conditions, and herpesvirus interactomes reached significance only in coronary artery disease (CMV) and type 2 diabetes (HHV8). Among other, non-herpes, viral interactomes, the one of HIV was MS-associated in the GWAS 2011, METACHIP1 and METACHIP2 but not in GWAS 2013. Other viral interactomes showing associations with MS were HBV (GWAS 2011), polyomavirus (GWAS 2013) and VIRORF (GWAS 2013). Other occasional associations with MS in the non-viral interactomes were AHR, Chlamydia and SIRT7 in the GWAS 2013 and AIRE in the GWAS 2011. Taken together, these results point to a disease-specific influence of EBV and, possibly, other herpesviruses on the development of MS.

### Common regulatory mechanisms exist among all the MS-associated interactomes

To better understand the overall picture that may emerge from the above results we evaluated the possibility that common regulatory mechanisms exist among all the MS-associated interactomes. To verify this hypothesis, we considered all the SNPs and genes that were responsible for the statistical enrichments of association of interactomes with MS. Specifically, we obtained n=10,788 nominally significant SNPs resulting from ALIGATOR analysis and the corresponding 741 MS-associated interactome genes (MS-AIG) (Figure 2A, Table S1). We mapped the SNPs and related MS-AIG in the human genome, showing a wide distribution on autosomal chromosomes (Figure 2B) and a prevalent localization in intronic and upstream/downstream regions (Figure 2C). This result is in line with the concept that genetic variants associated to multifactorial diseases affect the regulation of gene expression (Farh et al. 2015; Freund et al. 2018), reinforcing the plausibility of our observations. We then used RegulomeDB database (Boyle et al. 2012) to look for proteins whose binding close to SNPs extracted from ALIGATOR analysis is enriched. We overlapped the SNPs with the protein binding map generated by Encyclopedia of DNA Elements (ENCODE) and RoadMap Epigenomics (Boyle et al. 2012), obtaining a list of enriched proteins. RNA Polymerase II Subunit A (POLR2A) and CCCTC-Binding Factor (CTCF) displayed the highest regulatory potential (Figure 2D). Since each SNP can be bound by multiple proteins, we carried out an analysis of “co-occurrence” for POLR2A and CTCF evaluated using Fisher’s exact test. After controlling P-values for multiple-testing (FDR, Benjamini-Hochberg), co-occurrence of POLR2A and CTCF was significant (q<0.05). This is in line with data showing that SNPs in CTCF binding sites could alter chromatin topology and affect gene expression in the presence of RNA polymerase II (Tang et al. 2015).

**Figure 2:**
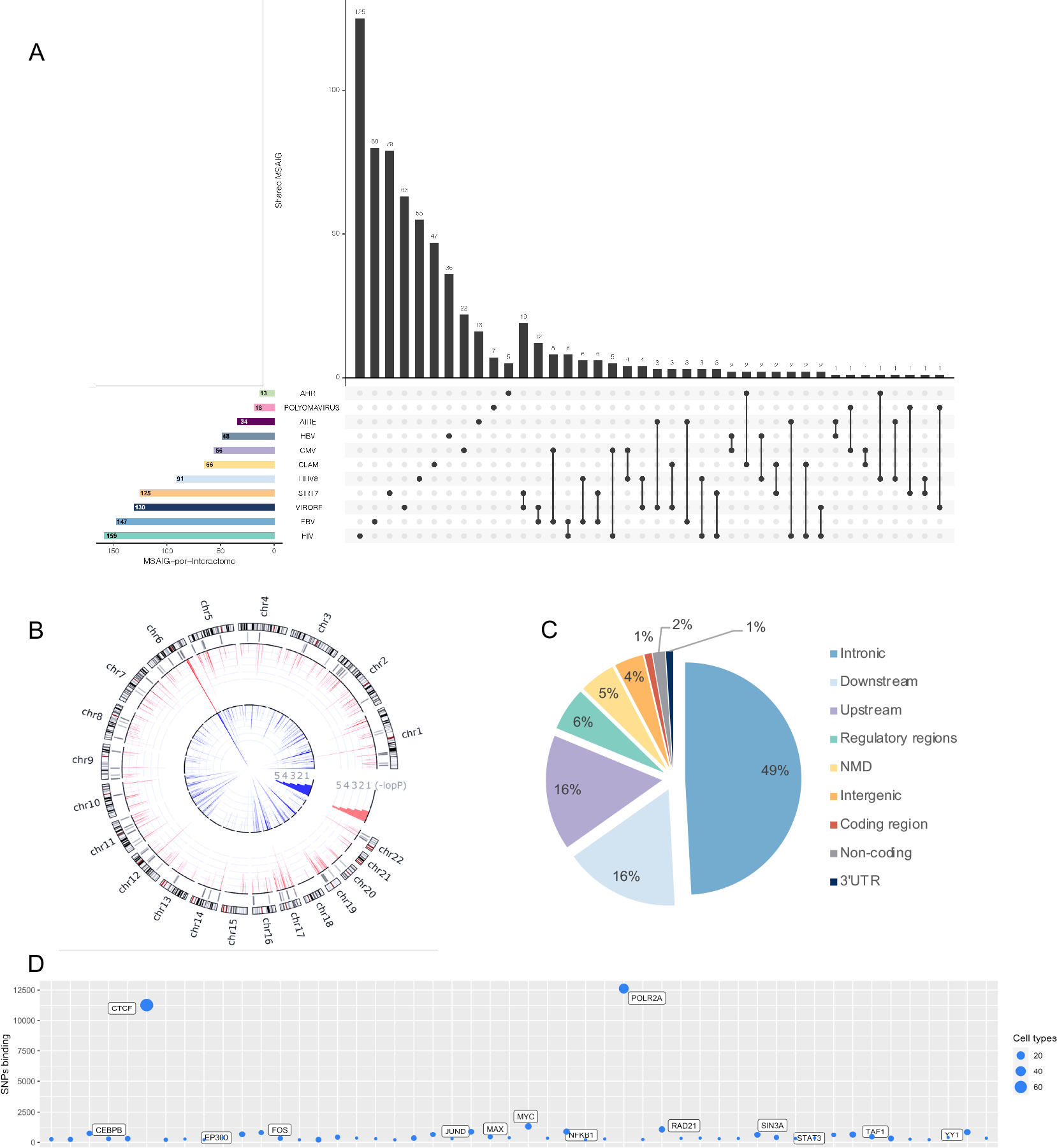
Genomic distribution of the candidate interactome SNPs and protein binding enrichment near MS-associated SNPs. (A) The UpsetPlot displays the number of MSAIG for each interactome of origin (x axis) and the intersections between gene sets with the respective size (y axis) (see also Table S1). (B) The circos plot shows the genomic distribution of the SNPs obtained from the candidate interactome analysis. The outer circle shows the chromosomes with related cytobands. The underlying grey lines indicate the physical position of the corresponding 741 MS-AIG (each line may represent up to 25 genes). The red and blue lines in the inner circles indicate the -log(Pvalue) for each SNP respectively from GWAS 2011 and GWAS 2013 (scaled from 0-20, which truncates the signal in several regions). (C) The pie chart represents the distribution of SNPs in different regions of the human genome, showing an enrichment in intronic and downstream/upstream regions. (D) The bubble plot displays the number of SNPs bound by a given protein obtained from different cells types (vertical axis) and the top 50 proteins enriched on MS-associated SNPs (horizontal bar). NMD = Nonsense-mediated mRNA decay; 3’UTR = three prime untranslated region.

### Dysregulations of MS-associated EBV interactors in blood and CNS transcriptomes

To evaluate the functional implications of the above results, we matched the 741 MS-AIG with microarray gene expression data obtained from peripheral blood leucocytes of 40 healthy subjects and persons with different MS phenotypes, including clinically isolated syndrome (CIS, n=46), relapsing-remitting (RR, n=52), secondary progressive (SP, n=21) and primary progressive (PP, n=23) MS (Srinivasan et al. 2017b). We calculated whether the MS-AIG were significantly enriched in the lists of genes differentially expressed in each MS phenotype. We took into consideration differentially expressed genes with p<0.05 and fold change >1.5. Totally, 464 MS-AIG were differentially expressed in at least one MS phenotype but no significant enrichment appeared in any of the 4 disease subtypes when compared with a random selection of transcripts from the whole transcriptome. However, when considering separately the MS-AIG from single interactomes, we observed an enrichment of differentially expressed MS-AIG from AIRE interactome in CIS patients, from AIRE and SIRT7 interactomes in RRMS, and from EBV interactome in PPMS (Table 1).

**Table 1:** MS-AIG differentially expressed in peripheral blood leucocytes. The enrichment of MS-AIG was calculated considering separately each interactome in distinct MS phenotypes. The associations reaching significance are highlighted in bold.

| MS phenotypes | N° of DE MS-AIG | P-value | OR | Frequency of MS-AIG in blood DEG (%) |  |  |  |  |  |  |  |  |  |  |  |
| --- | --- | --- | --- | --- | --- | --- | --- | --- | --- | --- | --- | --- | --- | --- | --- |
|  |  |  |  | AHR | AIRE | CHLAMYDIA | CMV | EBV | HBV | HHV8 | HIV | POLYOMAVIRUS | SIRT7 | VIRORF | GLOBAL TRANSCRIPTOME |
| CIS | 166 | 0.59 | 0.28 | 25.00 | **50** | 29.03 | 34.62 | 24.32 | 16.00 | 22.09 | 21.43 | 45.45 | 27.93 | 30.51 | 25.56 |
| RRMS | 170 | 0.34 | 0.89 | 25.00 | **40.63** | 25.81 | 19.23 | 22.97 | 24.00 | 27.27 | 21.90 | 45.45 | **34.23** | 27.97 | 23.66 |
| PPMS | 249 | 0.082 | 3.02 | 25.00 | 40.63 | 38.71 | 38.46 | **44.59** | 46.00 | 42.86 | 32.86 | 27.27 | 31.53 | 34.75 | 33.83 |
| SPMS | 226 | 0.53 | 0.37 | 31.25 | 37.50 | 27.42 | 32.69 | 33.11 | 28.00 | 37.66 | 32.86 | 9.09 | 30.63 | 34.75 | 32.38 |

CIS = clinically isolated syndrome; RR-MS = Relapsing-Remitting multiple sclerosis; PP-MS = Primary progressive multiple sclerosis; SP-MS = Secondary progressive multiple sclerosis; DEG= differentially expressed genes.

To understand the functional implications of our findings also at the central nervous system (CNS) level, we matched the 741 MS-AIG with microarray gene expression data obtained from post-mortem brain tissue of 20 people with secondary progressive MS (SPMS) and 10 non-neurological controls. Grey matter lesions (GML) and normal appearing grey matter (NAGM), from MS brains that exhibited the presence (F+) or absence (F-) of lymphoid follicles, were considered (Magliozzi et al. 2019). Again, we calculated whether the overall frequency of MS-AIG was significantly higher in the dysregulated gene lists (p<0.05 and fold change >1.5) than its expected frequency in a random selection of transcripts from the whole transcriptome. Overall, 169 MS-AIG were differentially expressed in at least one of the four conditions analyzed (GML-F+, GML-F-, NAGM-F+, NAGM-F-). The result was significant (Table 2). Considering separately the MS-AIG obtained from each MS-associated interactome, we observed a significant enrichment only for genes belonging to the EBV interactome in all the four conditions analyzed.

**Table 2:**
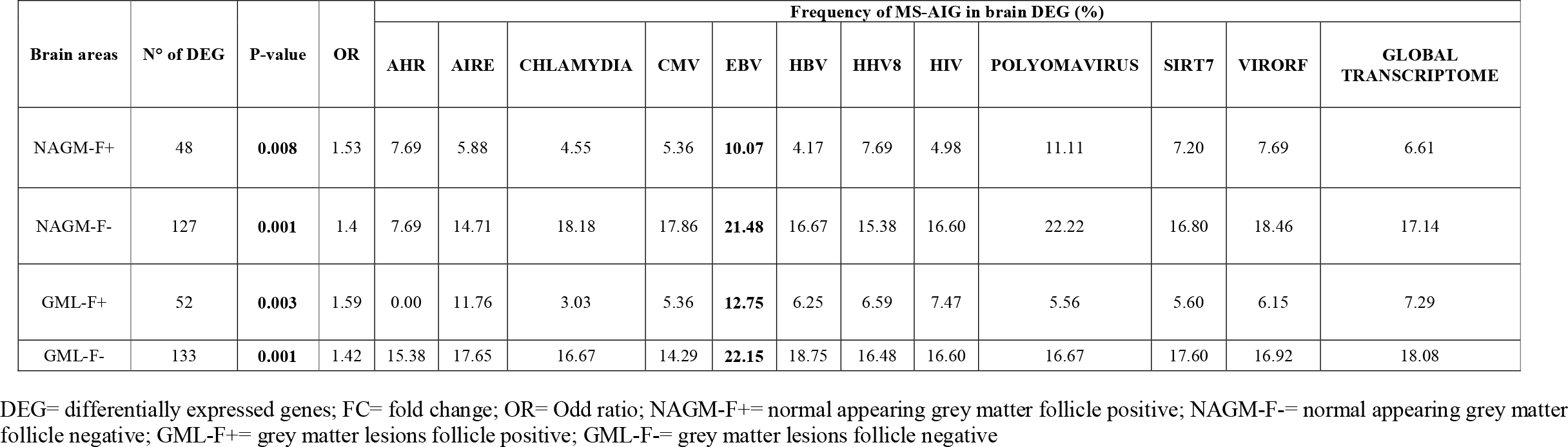
MS-AIG differentially expressed in brain. The frequency of MS-AIG was calculated in DEG of 4 post-mortem brain areas from secondary progressive MS. The enrichment of MS-AIG was also calculated considering separately each interactome in distinct brain areas. The associations reaching significance are highlighted in bold.

The above results from peripheral blood and CNS transcriptomes imply a potential role for EBV-related neuroimmunological response in the neurodegenerative and neuroinflammatory aspects of the disease. This prompted us to consider recent data showing an enrichment of EBV interactors in protein modules involved in Hereditary Spastic Paraplegia (HSP), an inherited degenerative axonopathy (Bis-Brewer et al. 2019) sharing clinical similarities with progressive MS. Importantly, it has been shown that persons with PPMS are enriched for HSP-related mutations (Jia et al. 2018). We therefore matched the 741 MS-AIG with the genes of the HSP-protein modules (n=295) and found 71 genes in common. Of these, 34 were included in EBV interactome (n= 147 from MS-AIG) (Figure S4). This number resulted larger than randomly expected (p-value<0.00001). In summary, gene expression data support the results of the candidate interactome analysis and show that the functional implications may be more condensed at the CNS level.

### Interactome-driven pathophysiology converges on the CD40 pathway in MS

To highlight the potential biological functions affected by the MS-AIG, we performed a pathway analysis using MetaCore^©^. The 741 MS-AIG were mapped to the internal MetaCore KnowledgeBase and a classification in canonical pathways was obtained. This analysis showed that the most critical biological process involved in MS through gene-environment interactions was CD40 signaling, followed by stress-induced antiviral cell response, oxidative stress and apoptosis (Figure 3A). This result was in agreement with studies showing the role of the CD40 pathway and that of B cells in MS pathophysiology (Afrasiabi et al. 2019; Field et al. 2015; Milo 2019; Smets et al. 2018).

**Figure 3:**
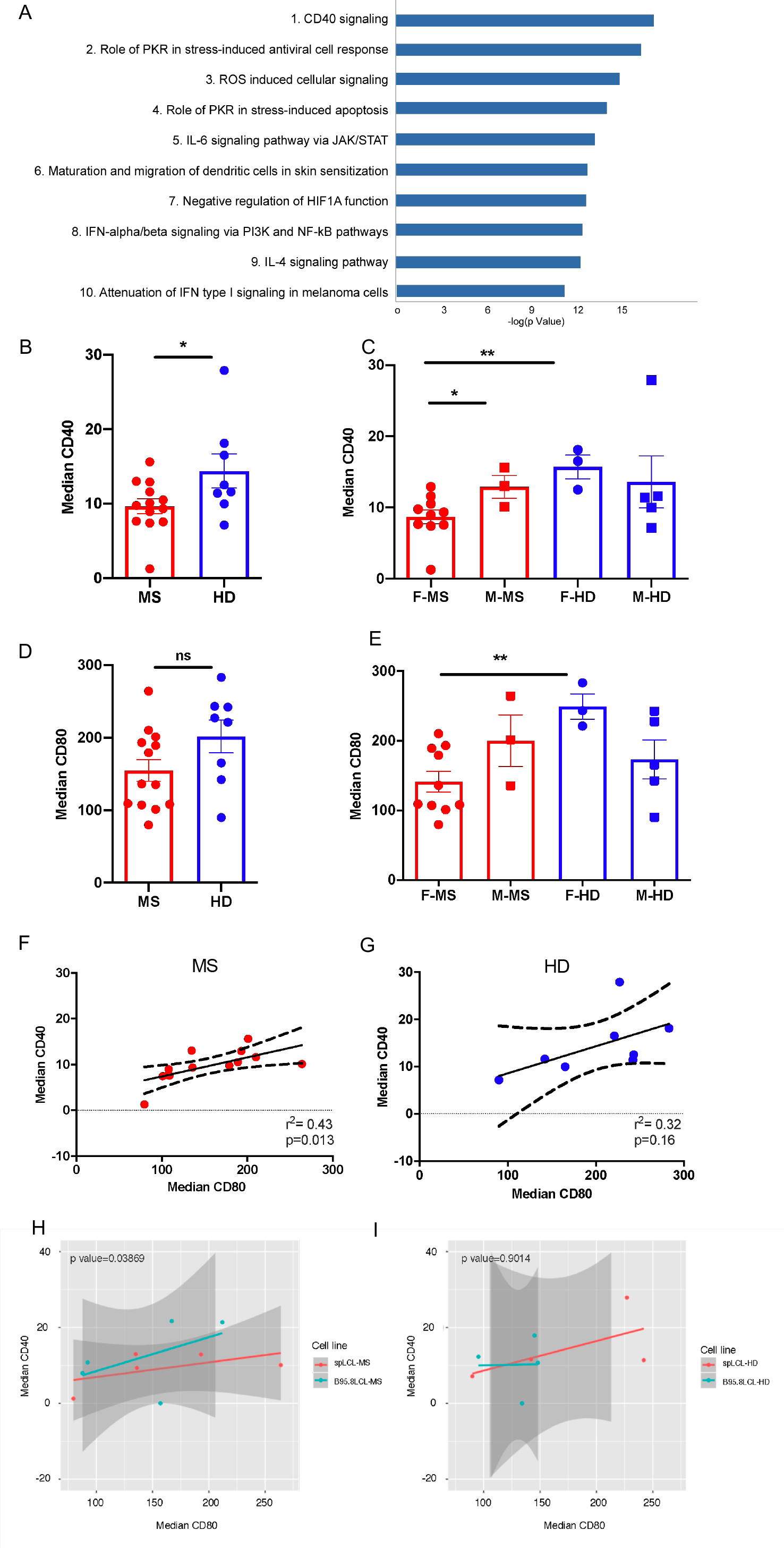
CD40 signaling analysis. (A) The top 10 pathways obtained from the analysis of the MS-AIG. The pathway analysis was performed using MetaCore and its canonical pathway maps. The lower p-value (-log Pvalue) means higher relevance of the genes within the pathway datasets. The CD40 signaling reached the highest relevance. (B) The CD40 protein level analyzed in MS-spLCLs (n=13) and HD-spLCLs (n=8) is downregulated in MS-spLCLs at unpaired T-test (*p=0.04). (C) The CD40 protein level stratified according to the gender, CD40 protein is under-expressed in females of MS group (F-MS) compared to the females of control group (F-HD;**p=0.006). (D) The CD80 protein level is comparable between MS-spLCLs and HD-spLCLs. (E) The CD80 protein level stratified according to the gender, CD80 protein is under-expressed in F-MS compared to the F-HD (**p=0.0037). (F-G) A correlation between CD40 and CD80 protein levels was evaluated by a simple linear regression analysis (Spearman correlation) in MS (r^2^= 0,43 p= 0,016; F) and in HD (r^2^= 0,32 p= 0,16; G). Dashed curves represent the confidence interval, the straight lines represent the correlation slope. The protein levels are represented as median fluorescence intensity. Data analysis was performed using Unpaired t-test F= female; M= male. (H-I) The effect of the EBV strain (spLCL vs B95.8LCL) and CD80 levels on CD40 protein expression in MS patients (n=5; H) and controls (n=4; I) were analyzed by a mixed-effect multiple linear regression model, setting subjects as random effect, and cell line context and CD80 levels as fixed effects.

To investigate whether CD40 alterations in MS could be driven by the interaction between host- and virus-specific factors we used flow cytometry to evaluate the surface protein expression of CD40 and co-stimulatory molecules CD80 and CD86 [known to be induced by CD40 and by EBV infection (Morandi et al. 2017)] in spontaneously outgrowing lymphoblastoid cell lines (spLCLs) obtained from 13 persons with MS (MS-spLCLs) and 8 healthy donors (HD-spLCLs). SpLCLs were obtained from B cells infected by the endogenous EBV strain: this condition allowed us to study the immunologic profile in a setting that may be more representative of a putative pathophysiological context where host and virus genetic variants interact in vivo. Results were compared with those obtained in B cells infected with the recombinant EBV strain B95.8 (B95.8LCL).

We found a reduction of CD40 protein level in MS-spLCLs, compared to HD-spLCLs (p= 0.04; Figure 3B). This was especially evident when we stratified the data according to sex: we observed a more pronounced difference in females with MS compared to healthy females (p=0.006), and a near-significant difference (p=0.06) when we compared females and males in the MS group (Figure 3C).

Analyzing the CD80 protein level, we found a slight reduction of the surface protein in MS-spLCLs compared to HD-spLCLs (p= 0.089; Figure 3D); once again a significant difference emerged when we compared the affected and healthy female groups (p= 0,0037, figure 3E). The CD86 protein levels showed no differences between MS-spLCLs and HD-spLCLs (Figure S5). We then performed a simple linear regression analysis to evaluate a possible interdependence between CD40 and CD80 protein levels. The two protein levels were correlated in MS-spLCLs (r^2^= 0,43 p= 0,016; Figure 3F) but not in HD-spLCLs (r^2^= 0,32 p= 0,16; Figure 3G).

The MS associated SNPs rs4810485, which maps within the CD40 gene, and rs9282641, which maps within the CD86 gene, have been reported to be associated with a reduced CD40 expression and an increased expression of CD86 on the B cell surface (Smets et al. 2018). We did not observe this correlation in the spLCLs (Figure S6). We also measured CD40, CD80 and CD86 protein expression in B cells infected with recombinant EBV strain B95.8 (B95.8LCL) in 6 patients and 6 HD but did not find significant differences (Figure S7).

Finally, we performed a multiple linear regression to assess the possible influence of the EBV strain (spLCL vs B95.8LCL) on CD40 expression level controlling for CD80, since CD80 levels were correlated with CD40 protein expression (see Figure 3F). For this analysis, we considered only spLCL and B95.8LCL cell lines obtained from the same subject, at the same time point. This experimental setting allowed us to observe a lower CD40 expression in MS-spLCLs compared to MS-B95.8LCLs (p=0.04; Figure 3H-I), suggesting an effect of MS-associated EBV variants on CD40 protein level. With the possible limitation of the reduced sample size of spLCLs under scrutiny, we were not able to link this effect to MS-associated EBV variants previously found in the EBNA2 gene (Mechelli et al. 2015). This issue deserves further investigation, by increasing the number of spLCLs scrutinized and extending the analysis to other EBV genes.

We then used MetaCore^©^ to identify the biological functions possibly affected by the MS-AIG dysregulated in the CNS. Once again, we found CD40 signaling as the pathway that more clearly emerged from the disease-predisposing, host-environment interaction in MS, also at the lesion level. By using digital droplet-PCR (ddPCR) we analyzed the CD40 mRNA level in the same RNA samples obtained from cortical lesions of post-mortem progressive MS cases used for the microarray gene expression analysis. CD40 expression was increased in the subpial cortical lesions of MS cases compared to the normal grey matter of non-neurological controls (fold change: 1,9, p= 0.007; Figure 4A, Table S2). By using immunohistochemistry, the CD40 expression was observed mainly in active lesions, either in the grey matter (Figure 4B, C) or in the white matter (Figure 4E, F). Sporadic CD40+ cells were also observed in normal appearing grey matter (Figure 4D) and normal appearing white matter (Figure 4G), while CD40 expression was not detected in inactive lesions and in non-neurological controls. In particular, CD40 expression was detected in cells morphologically resembling activated microglia, as further supported by double immunolocalization with the MHC class II marker (Figure 4 insets in C and in F). Though the experimental settings (spLCLs and brain lesions) do not allow to conduct analyses on large numbers, the results suggest that the interaction between host- and EBV-specific factors may lead to CD40 signaling alterations.

**Figure 4:**
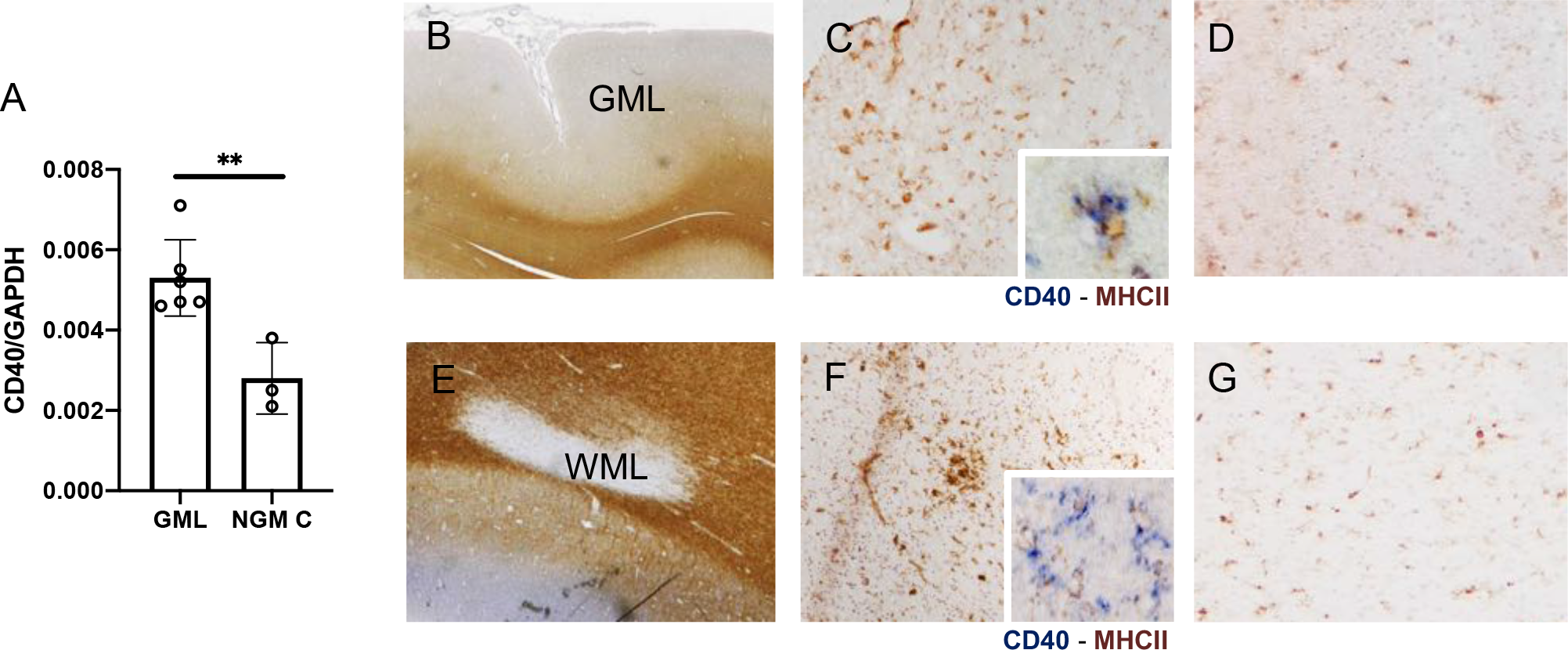
CD40 analysis in post-mortem progressive MS brain tissues. (A) ddPCR analysis of CD40 gene expression, referred to the housekeeping gene GAPDH, in 6 GML and in 3 NGM from controls. The CD40 mRNA level is up-regulated in diseased tissue at Unpaired t-test. Data are shown as mean ± SD. **p= 0.007. (B-G) Neuropathologycal assessment of the CD40 protein expression in post-mortem MS cases. CD40 protein expression was observed mainly in active lesions, either in the grey matter (B, C) or in the white matter (D, F), in cells morphologically resembling activated microglia, as further supported by double immunolocalization with the MHC class-II marker of activated microglia/macrophages (insets in C and in F). Sporadic CD40+ cells were also observed in normal appearing grey matter, NAGM (D) and normal appearing white matter, NAWM (G). Original magnifications: 10x (B, D); 100x (C, D, F, G); 400x (insets in C and in F).

### EBV interacting proteins have the highest therapeutic potential in a Priority Index analysis of MS-AIG

A recent genetics-led drug target prioritization approach (Fang et al. 2019) represented an interesting resource to translate our results into therapeutically relevant information. Fang and colleagues developed a Priority Index (Pi) pipeline, based on GWAS variants for several immunopathological traits, including MS, that allowed us to investigate whether, and to what extent, the MS-AIG may contribute to the discovery of new druggable targets. Among the first 150 positions in Pi Ranking for MS, 26 appeared in MS-AIG (Figure 5A). This enrichment was specifically due to 13 genes belonging to the EBV interactome (p =0.0045). We also matched the MS-AIG with the list of genes showing the highest druggability among the top 1% genes prioritized in the work by Fang et al. Interestingly, 24 out of the previously identified 26 MS-AIG were included in high-Pi rated genes and were highly-druggable targets. Then we looked at the position of the 741 MS-AIG genes within the actual landscape of drug target genes (including Phase 1 to Phase 4 trials) in immunological diseases. Six of these genes proved to already be the object of clinical investigation for MS, while 10 were not specifically intended to treat MS, but are involved in the treatment of other diseases (Figure 5B and Table S2). Among these were the highest Pi-prioritized MS-AIG, such as IL6, JAK1 (HBV interactome), JAK3 and SYK (EBV interactome). The results therefore suggest that a pharmaceutical targeting of relevant gene-environment interactions is a feasible strategy that may potentially advance the efficacy of MS treatments.

**Figure 5:**
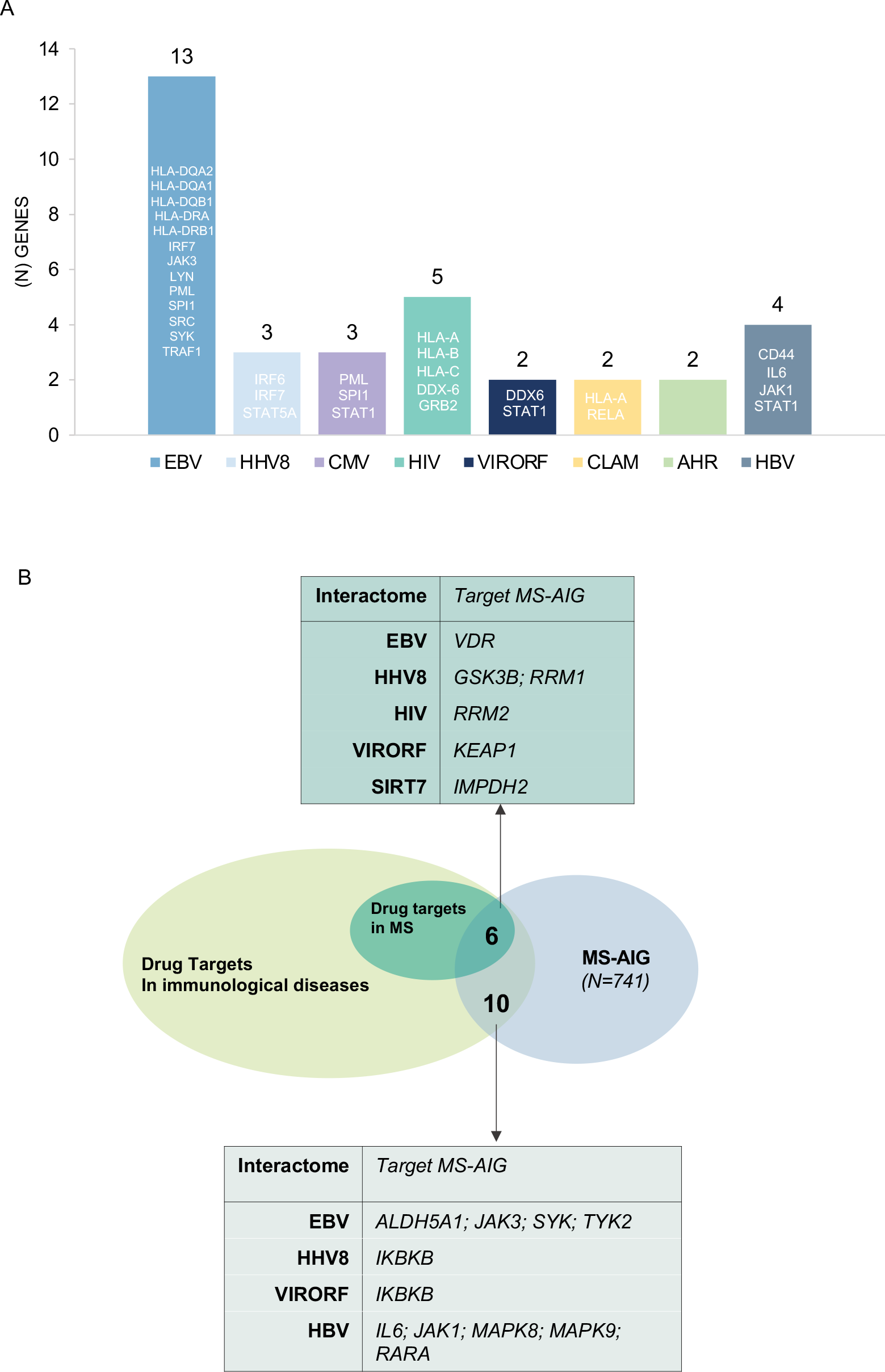
MS-AIG Pi-Prioritization and distribution in actual drug targets for immunological diseases. (A) The figure displays MS-AIG present in the top 150 Pi-prioritized genes for MS (see text, and Fang et al. 2019). *Y axis* shows the number of genes and their identifier, *x axis* shows the corresponding interactomes. (B) The diagram shows the overlap between the 741 MS-AIG and the actual clinical proof-of-concept drug targets for immunological diseases, including MS, as resulting from Fang et la, 2019. The 16 overlapping genes are split between MS and other immunopathological conditions and reported in the charts with the corresponding interactome.

## DISCUSSION

The association between MS and EBV is a long recognized one (Ascherio and Munger 2016). Starting from the indications of a preliminary study (Mechelli et al. 2013), we here show that the interaction between genotype and herpesviruses may have causal implications in MS, in ameaningful setting that takes into account new and more accurate data and additional in silico and in vitro analyses. The results now indicate that this finding is specific for MS and not for other immune-mediated diseases. Peripheral blood and CNS transcriptome data in MS show a dysregulation of MS-associated EBV interactors in both compartments, further validating the result (Gustafsson et al. 2014; Lamparter et al. 2016). Pathway analyses of the MS-AIG and protein expression experiments in spLCLs from persons with MS and controls, show that the interaction between MS-AIG and viral exposures may converge towards the dysregulation of the costimulatory receptor CD40 and its pathway. In accord with the observed transcriptomic dysregulations in MS lesions, matching MS-associated interactome genes with protein interaction modules from inherited axonopathy-causing genes, supported a link between MS-associated EBV interactors and neurodegeneration. Finally, we show that products of the MS-AIG may be prioritized as drug targets.

Compared to our previous results, the new “candidate interactome” analysis confirms the relevance of EBV, shows the emergence of other herpesviruses and appears to disconfirm other viruses (e.g. HBV) that were significant in the previous study. These results are in line with the sero-epidemiological literature, with obvious biological similarities between EBV and other herpesviruses, and with biological plausibility. Furthermore, the results indicate that this kind of analyses become increasingly meaningful and reliable as new genetic and protein-protein interaction data become available (Huttlin et al. 2021; Wang et al. 2021). It is remarkable that, in a recent study considering a multi-omics approach in MS (Badam et al. 2021), the analysis identified a module of 220 genes strongly associated with risk factors in MS. This module shows a significant overlap with our 741 MS-AIG and with the EBV interactome genes whose expression we found dysregulated in the transcriptome analyses (Figure S8). Concerning other viruses, and again in comparison with our previous work, HIV retained a significant association. In spite of the lack of epidemiological links between HIV and MS, the result will deserve further attention as HIV can induce demyelination (Berger et al. 1989; González-Duarte et al. 2011; Johnson 1994). Furthermore, HIV may share some functional kinship with the HERV family of endogenous retroviruses potentially involved in the pathogenesis of MS and of other neurodegenerative diseases (An et al. 2001; Küry et al. 2018). Finally, our finding of CTCF acting as a common regulator of the MS-AIG reinforces the idea that the MS-AIG are disease-relevant genes that may be affected by the regulatory influence of EBV. In fact, it has been shown that CTCF-bound sites show enhancer-blocking activity towards developmental genes associated with human diseases (Martin et al. 2011) and overlap with sites where EBV alters accessible chromatin (Lamontagne et al. 2021), subverting the host’s gene expression program (Ersing et al. 2017; Zhou et al. 2015).

Support to our results comes also from two recent studies on low-frequency and rare-coding variants associated with MS (Consortium 2018; Vilariñ o-Güell et al. 2019). Altogether, these studies identified eighteen genes harboring MS-associated low-frequency coding variants. Surprisingly, half of these are either EBV interactors or implicated in EBV-related pathophysiologies (for details see Table S1). In a third study (Vidmar et al. 2019), MS-associated rare variants could be linked to molecular mechanisms connecting EBV with interferon-beta signaling and autophagy. Finally, the SNP with the highest association with white matter lesion volume in the recent genome-wide association study of brain magnetic resonance imaging phenotypes of the UK Biobank (Elliott et al. 2018), is in the 5’ UTR of EFEMP1, a gene that codes for another EBV interactor (its product interacts with two EBV proteins, BFLF2 and BRRF1) (Gulbahce et al. 2012).

Autoimmune diseases have common and disease-specific signatures, recently highlighted by gene expression studies of target tissues (Szymczak et al. 2021). In the present study, we applied the interactome analysis to immune-mediated diseases other than MS, where epidemiological studies have found associations with EBV, though probably less robust compared to MS. In these conditions, our data do not support the idea that the interaction between EBV and the genetic background of the host may contribute to disease etiology. The difference between MS and the other immune-mediated conditions appears plausible in the light of differences in pathophysiology and response to treatments. Distinctions have been proposed in genetic (Gregory et al. 2012) and immunological studies on the tumor necrosis factor (TNF) system (Gao et al. 2017; Veroni et al. 2010), one of the most ancient antiviral mechanisms (tenOever 2016). In addition to suggestions coming from mechanistic studies, clinical evidence is clear cut: it involves not only the well-known, opposite effects of anti-TNF therapies (highly beneficial in RA and detrimental in MS) but also immunotherapies that exploit cytokines with antiviral actions, such as interferon-beta. These are effective in MS but not in RA (Genovese et al. 2004; van Holten et al. 2005) or in other immune-mediated diseases where they may concur in the development of the diseases (Axtell et al. 2011). Furthermore, the response to anti-B lymphocyte therapies in MS, including in the progressive form of the disease, may be indicative of a role of EBV in disease pathophysiology as this virus remains for the lifetime of the host in memory B cells. Of note, in RA, anti-B lymphocyte therapies may not be as effective as anti-TNF treatments (González-Vacarezza et al. 2014).

Despite these differences, other mechanisms of EBV involvement in autoimmunity cannot be excluded. For example, the binding of EBNA2 to genomic intervals associated with various autoimmune diseases, including MS, suggests another mechanism that operates across diseases (Harley et al. 2018; Ricigliano et al. 2015). Furthermore, also in view of results about HHV-6 and HHV-7 in Alzheimer’s disease and in a non-human primate model of MS (Leibovitch et al. 2018; Readhead et al. 2018), it will be necessary to study more immune-mediated diseases and various other “candidate interactomes” of viral origin to draw a complete picture of the long-suspected role of viruses in autoimmunity and in neurodegeneration. Incidentally, given the pleiotropic effects of EBV on the immune response, the negative result in immune-mediated diseases other than MS to some extent controls for the possibility that the association between EBV interactome and MS derives from effects of the genetic variants on the outcome (MS) that are not exclusively mediated by the exposure (EBV) (horizontal pleiotropy) (Davey Smith and Hemani 2014; Davies et al. 2018).

An elegant study comparing gene expression in unselected peripheral blood B cells and B95.8 infected B cells in healthy individuals recently showed that EBV infection affects MS risk genes (Afrasiabi et al. 2019). We now extend this result to the disease state: the analysis of peripheral blood and brain cortex transcriptome data confirmed that, among the MS-AIG, those that are part of the EBV interactome are more frequently dysregulated than expected. Of the eight sets of gene expression data (four from the peripheral blood and four from brain cortical areas), MS-AIG belonging to the EBV interactome were significantly dysregulated in five. The dysregulation was more consistent in transcriptomes from the four brain tissue types (4/4) than in those from the peripheral blood (1/4). This result, particularly in the datasets from cortical tissue devoid of lymphoid follicles, may be to some extent unexpected given the propensity of EBV to infect B lymphocytes and influence the immune response. However, astrocytes and microglia activation, which have a key role in MS inflammatory response (James et al. 2020) in MS, may express the EBV receptor CD21 (Lindblom et al. 2015) and may be infected by the virus in this disorder (Hassani et al. 2018), suggesting that EBV may entertain pathogenic interactions also outside the B cell compartment. Analyses of the most recent MS-GWAS (Consortium 2019b), which shows an enrichment for MS genes in microglia, may clarify this issue, though it is already intriguing that the MS variant with the strongest eQTL in microglia, CLECL1, is an EBNA3C-regulated gene in lymphoblastoid cell lines (Zhao et al. 2011) and is upregulated in tonsillar B cells after CD40 ligation (Ryan et al. 2002). Here we show that EBV interactors within the MS-AIG are enriched in protein interaction modules derived from genes that cause hereditary spastic paraplegia (Bis-Brewer et al. 2019). In this disease, that has no clear evidence of an immune-mediated component but shares neurodegenerative features and clinical similarities with MS, a potential link with EBV infection (Bis-Brewer et al. 2019) and with MS (Jia et al. 2018) had been already suggested. Overall, these data imply that the contribution of EBV to CNS damage may, to some extent, be independent of a canonical inflammatory context. Whether or not the hypothesis of EBV acting at the CNS level be supported by its presence in latently infected B cells at the lesion level or in meningeal lymphoid follicles remains a controversial issue (Aloisi et al. 2010; Lassmann et al. 2011; Ramesh et al. 2020; Sargsyan et al. 2010; Serafini et al. 2007; Willis et al. 2009).

The CD40 pathway (a key factor in antigen presentation, immune regulation, B lymphocyte development, germinal center formation, somatic hypermutation and class-switch recombination) emerges among the biological functions affected by the MS-AIG, again in accord with data comparing peripheral blood and B95.8 infected B cells (Afrasiabi et al. 2019). This result is coherent with what is known about the role of the CD40 molecule in MS, where the risk allele affects the membrane-bound molecule and splice forms of decoy receptors (Field et al. 2015; Smets et al. 2018), once again in an opposite way between MS and other immune-mediated diseases such as rheumatoid arthritis (Raychaudhuri et al. 2008), systemic lupus erythematosus (Vazgiourakis et al. 2011), autoimmune thyroiditis (Tomer et al. 2002) and Kawasaki disease (Onouchi et al. 2012). With respect to the CD40 expression in LCLs, the data suggest an effect of MS-associated EBV variants on the downregulation of the membrane-bound molecule. However, due to the small numbers of spLCLs available, we were unable to understand which EBV variants are responsible for this effect. In more general terms the results are in line with the idea that viruses and disease risk alleles may induce or enhance diseases through similar mechanisms (Chen and Xia 2019; Gulbahce et al. 2012) and with the observation that the EBV-induced regulatory perturbations are often in the direction of a reduced gene expression (Zhou et al. 2015). Here, we suggest also a broader involvement of the CD40 signaling pathway and outline the gene-environment interactions that may generate its involvement in MS. This information is relevant as the CD40 pathway may participate in the creation of memory B cells (Baker et al. 2018) and in the activation of a repertoire of autoreactive, brain-homing CD4+ T cells in a way that does not seem to directly depend on the CD40 molecule itself (Jelcic et al. 2018). Interestingly, EBV-specific CD4+ T cells that cross-react with myelin and neuronal epitopes are part of this repertoire (Wang et al. 2020), suggesting the possibility that EBV participates in MS pathogenesis through multiple and synergic mechanisms.

The fact that we found increased CD40 expression by microglia/macrophages in active grey matter and white matter lesions of progressive MS cases may support the hypothesis that this molecule contributes to the active phase of lesion pathology and may therefore represent a potential candidate target for future therapeutic strategies (Jäckle et al. 2020). Previous experimental autoimmune encephalomyelitis (EAE) studies demonstrated that CD40 is necessary for the activation of microglia cells since the initial disease stages (Gerritse et al. 1996; Ponomarev et al. 2006). Our findings support and expand these data in the context of the human disease: the increased CD40 expression in NAGM and NAWM (even in absence of demyelination) in cell resembling activated microglia points to a pivotal role of brain resident innate immunity in disease initiation. CD40 may thus contribute to bridge the innate and adaptive immune response in neuroinflammation, deserving further and more in-depth analysis, given its clinical implications. In this context, our approach offers the opportunity to pinpoint therapeutic targets, prioritized on a genetic basis (Fang et al. 2019), that are related to “environmental” exposures. As such, these targets are in principle more actionable than others based on an ongoing host pathophysiology, particularly for preventive purposes. The Pi analysis of the MS-AIG, once more, highlighted the EBV interactome as a relevant reservoir of therapeutic targets and exposed MS-specific target pathways (e.g. interferon signaling) among which CD40 ranked highest.

In accord with recent suggestions about the importance of understanding genetic variability at the systems level rather than in individual genes (Li et al. 2019) this study indicates that the interaction between host genotype and EBV (including other Herpesviruses) is relevant for MS etiology, reinforcing the rationale for future studies that attempt to decipher disease mechanisms by looking at virus-mediated pathology (Müller-Schiffmann et al. 2021). It also strengthens the rationale for experimental medicine approaches aimed at advancing therapies while proving the causality of viral agents in MS. The results of a recent study on EBV-specific adoptive T cell therapy in patients with progressive MS support this perspective (Pender et al. 2018).

## Supporting information

Supplemental informations_Mechelli et al. 2021

## Acknowledgments

This work was supported by the Italian Multiple Sclerosis Foundation (FISM) (grant number 2014/R/12 to RM). CENTERS is a Special Project of the Italian Multiple Sclerosis Foundation (FISM).

G.M. is funded by Fondazione Italiana Sclerosi Multipla (FISM; grants 2016/R/18 and 2018/S/5), Progetti di Rilevante Interesse Nazionale (PRIN; grant 2017K55HLC 001), and Ministero della Salute (grant RF-2019-12371111). P.d.C. is supported by FISM (grant number 2018/R/4).

## Author contributions

R.M., R.U., G.R., M.S. conceived and designed the study. R.M. selected and constructed the candidate interactomes and performed digital droplet-PCR analysis. R.U., V.R., G.B. and R.U.P. performed the bioinformatics works. R.M., R.U., D.C., L.B., G.R., M.S. analyzed the data. S.S. and C.F. provided blood transcritptome data and were involved in data analysis. R.MA. and R.R. provided brain transcriptome data and were involved in data analysis. R.MA. performed neuropathological brain tissue analysis. D.F.A. and G.G. performed cytofluorimetric analysis. S.R. and G.B. conceived and drawn the graphical abstract. R.B., E.A. and P.T. provided lymphoblastoid cell lines and were involved in data analysis. A.F., M.F., MCB and S.R. provided reagents, analysis methods, and instrumentation. S.DA. carried out sample genotyping and was involved in data analysis. R.M., G.R. and M.S. provided data interpretation and wrote the manuscript. C.F., L.B., S.DA., P.D.C., G.M., A.U., D.D.S., P.M. and D.C. provided critical reading of the manuscript. IMSGC and WTTTC2 provided GWAS data.

## Competing interests

Mechelli R., Renato U., Rinaldi V., Bellucci G., Angelini D.F., Guerrera G., Bigi R., Fornasiero A., Ferraldeschi M., Romano S., Buscarinu MC., Srinivasan S, Anastasiadou E., Trivedi P., Di Silvestre D., Mauri P., De Candia P., Battistini L., D’Alfonso S., Magliozzi R. and Ristori G. declare no conflict of interest.

Centonze D. is an Advisory Board member of Almirall, Bayer Schering, Biogen, GW Pharmaceuticals, Merck Serono, Novartis, Roche, Sanofi-Genzyme, Teva and received honoraria for speaking or consultation fees from Almirall, Bayer Schering, Biogen, GW Pharmaceuticals, Merck Serono, Novartis, Roche, Sanofi-Genzyme, Teva. He is also the principal investigator in clinical trials for Bayer Schering, Biogen, Merck Serono, Mitsubishi, Novartis, Roche, Sanofi-Genzyme, Teva. His preclinical and clinical research was supported by grants from Bayer Schering, Biogen Idec,

Celgene, Merck Serono, Novartis, Roche, Sanofi-Genzyme e Teva

Pizzolato Umeton R. is supported by the National MS Society.

Farina C. received research support from Merck-Serono, Teva, Novartis.

Uccelli A. has received consulting honoraria including as part of advisory board and/or speaker fees from Sanofi Genzyme, Roche, Biogen, Novartis, TEVA and Merck.

Matarese G. reports receiving research grant support from Merck, Biogen, and Novartis and advisory board fees from Merck, Biogen, Novartis, and Roche

Reynolds R. received research support from the UK MS Society and MedImmune and has received speaker honoraria from Roche and Novartis.

Salvetti M. received research support and consulting fees from Biogen, Merck-Serono, Novartis, Roche, Sanofi, Teva

## Supplementary figure legends

**Figure S1**: **Schematic representation of METACHIP datasets construction, related to Figure 1**.

Metachip 1 and 2 were built as the position-wise union of GWAS and Immunochip. Each dataset is represented by a line in which sample SNPs are indicated with different colors depending on their source dataset (black round for SNPs coming from GWAS and empty round for SNPs coming from Immunochip). When a given SNP existed in both GWAS and Immunochip, we gave preference to GWAS in Metachip1 and to Immunohip in Metachip2.

**Figure S2: Spearman correlation in MS GWAS, related to Figure 1**.

Linear correlations between the size of single interactomes and their cumulative p-value of association with MS calculated for each GWAS dataset (GWAS 2011, GWAS 2013, METACHIP 1 and METACHIP 2). The 95% confidence intervals of the Spearman correlation are greyed out in all the analyses.

**Figure S3: Spearman correlation in non-MS GWAS, related to Figure 1**.

Linear correlations between the size of single interactomes and their cumulative p-value of association with complex diseases calculated for each GWAS dataset (Bipolar disorder, celiac disease, Chron disease, coronary artery disease, hypertension, rheumatoid arthritis, T1 diabetes, T2 diabetes). The 95% confidence intervals of the Spearman correlation are greyed out in all the analyses.

**Figure S4: Overlap between MS-AIG and genes of HSP-dominant expanded modules**.

(A) Overlap between MS-AIG and HSP-protein modules genes. Out of 71 shared genes, 34 resulted EBV interactors. (B) Null distribution of EBV interactors included in 100.000 random samples of the same size of the 71 shared genes, extracted from MS-AIG. The observed number of EBV interactors (N=34 out of 71 shared genes, red line) was larger than randomly expected (p value<0.00001).

**Figure S5: The protein expression of CD86 on spLCLs is comparable between MS (MS-spLCLs, n=10) and controls (HD-spLCLs, n=6; A), related to Figure 3**. The protein levels are represented as median fluorescence intensity. The data analysis was performed using Unpaired t-test.

Ns= not statistically significant.

**Figure S6: Genotype independent CD40 and CD86 protein expression on spLCLs, obtained from MS patients and controls (HD), related to Figure 3**. (A) The rs4810485*T is the CD40 MS risk allele. The protein expression levels (represented as median fluorescence intensity) are independent from the genotype when analyzed overall (GG, n=12; TG, n=7; TT, n=2) and when MS (GG, n=7; TG, n= 4; TT, n=2) and HD (GG, n=5; TG, n=3; TT =not detected) were analyzed separately.

(B) The rs9282641*G is the CD86 MS risk allele. The protein expression levels are independent from the genotype when analyzed overall (GG, n= 15; GA, n=2) and when MS (GG, n=10) and HD (GG, n= 5; GA, n=2) were analyzed separately.

The data analysis was performed using 1-way ANOVA, Bonferroni post-test. Ns= not statistically significant

**Figure S7: Protein expression of CD40, CD80 and CD86 on B95-8LCLs surface, related to Figure 3**.

The protein levels are comparable between MS (n=6) and HD (n=6)-derived B95-8LCLs.

On *Y-axis* are represented the median fluorescence intensity of each protein. The data analysis was performed using Unpaired t-test. Ns= not statistically significant.

**Figure S8: Overlap between MS-AIG and the MS disease module derived from multi-omics analysis by Badam et al, 2021**.

(A) Overlap between all 741 MS-AIG and the 220 genes forming the MS module defined by Badam et al, 2021 through comparative multi-omics analysis.

(B) Overlap between the MS Multi-Omic Module and the MS-AIG differentially expressed in the brain, blood and both. The overlap is considered significant if p <0,05 (Fisher’s exact test, calculated through geneOverlap package in R software). ns= not significant.

## STAR METHODS

### LEAD CONTACT AND MATERIALS AVAILABILITY

Further information and requests for resources and reagents should be directed to and will be fulfilled by the Lead Contact, Marco Salvetti. This study did not generate new unique reagents.

#### EXPERIMENTAL MODEL AND SUBJECT DETAILS

##### Lymphoblastoid cells lines generation and DNA extraction

Sp-LCLs, carrying endogenous EBV, were generated from 13 subjects with relapsing remitting MS (Polman et al. 2011) and 8 healthy donors matched for age, sex and geographic origins (Chiara et al. 2016). Briefly, peripheral blood mononuclear cells (PBMCs) obtained by density centrifugation over Ficoll–Hypaque were seeded in 96-flat well plates and cultured in RPMI1640 medium, supplemented with 20% FBS (HyClone), 1% L-Glutamine, 100 IU penicillin, and 100μg/ml streptomycin. Cyclosporin A (1 μg/ml, Calbiochem) was added to the medium to inhibit T-cell activation and cultures were fed twice a week.

PBMC infected with EBV B95.8 strain (B95.8-LCL) were generated from 5 relapsing remitting MS subjects and 4 healthy donors incubating for 2 hours at 37°C the primary cells in a mix containing: virus (10x concentrated), Cyclosporin A (1 µg/ml) and RPMI1640 medium supplemented with 20%FBS, 1% L-Glutamine, 100 IU penicillin and 100 µg/ml streptomycin. Cells were seeded in 24-well plates and fed twice a week.

Genomic DNA was extracted from five million cells using the QIAamp DNA mini kit (Qiagen, Venlo, the Netherlands).

All subjects with MS were sampled during the stable phase of the disease and were free from disease modifying therapies. The local institutional review board approved the study and all participating subjects gave written informed consent.

##### Flow cytometry analysis

The phenotypic characterization of LCLs obtained from MS patients and HD was performed through multiparametric flow cytometric (MPFC) analysis. The following monoclonal antibodies were used: anti-CD40 PE (Pharmingen), B7-1 (anti-CD80 PE-Cy7) (Beckman Coulter), B7-2 (anti-CD86 APC) (Miltenyi), anti-CD19 APC-eFluor780 (eBioscience) and LIVE/DEAD Fixable Aqua Dead Cell Stain (ThermoFisher). The staining was performed with 0.5⨯x10^6^ cells with antibodies at previously defined optimal concentrations and incubated for 30 minutes at 4°C in the dark. The samples were acquired on a CyAn flow cytometer (Beckman Coulter), equipped with three lasers and able to measure up to 15 parameters simultaneously on each cell. For each sample, approximately 300,000 cells were selected based on scatter parameters, physical size (FSC) and grain size (SSC), and the analysis was conducted after the exclusion of dead cells and coincident events. Data were compensated and analyzed using FlowJo v10.5 (TreeStar, Ashland, OR, USA).

Statistical analyses were performed with GraphPad Prism 5.0 program.

##### CD40 and CD86 SNPs genotyping

We genotyped rs4810485 SNP in CD40 gene and rs9282641 SNP in CD86 gene in 26 spLCLs, using PCR amplification followed by Sanger sequencing.

The following PCR primers were used:

CD40 rs4810485: primers: forward: 5’-TGGTGATCAGAGGGCTGGAGAAA -3’, reverse: 5’-TCTCCACTCCTACCACAAGGGC-3’

CD86 rs9282641: primers: forward5’-AGGGCTTTACACTCATGCTCCGAG-3’, reverse: 5’-TTTTACTCATTCTCTGGCCGGCCA-3’

PCR amplicons were analyzed for SNP genotypes by Sanger direct sequencing using the Big-Dye Terminator v3.1 sequencing kit (Applied Biosystems, Foster City, California, USA) on an automated 3130xl Genetic Analyzer (Applied Biosystems, Foster City, CA, USA).

PCR and sequencing conditions are available on request.

##### Candidate interactome analysis

The “candidate interactome” approach was applied as previously described (Mechelli et al. 2013). In brief, we obtained 20 candidate interactomes from the literature. Six were manually curated (EBV, HBV, CMV, HHV8, JCV, Inflammasome); 7 were obtained from databases of molecular interactions: AIRE, VDR, AHR, SIRT1, SIRT7 from BIOGRID (http://thebiogrid.org) (Chatr-Aryamontri et al. 2015), Polyomavirus from VirusMentha database (Calderone et al. 2015), Human-microRNA targets from miRecords database (Xiao et al. 2009); 7 were obtained from published high-throughput experimental approaches: VIRORF (Pichlmair et al. 2012), HIV (Jäger et al. 2011), HCV (de Chassey et al. 2008), h-IFN (Li et al. 2011), H1N1 (Shapira et al. 2009), HDAC (Joshi et al. 2013), Chlamydia (Mirrashidi et al. 2015) (Figure 1A).

The manually curated interactomes were obtained by selecting only those interactions that were reported by two independent sources or were confirmed by the same source with distinct experimental approaches. In all cases, we considered only physical-direct interactions (Mechelli et al. 2013). All SNPs which did not pass quality checks in the GWAS studies were filtered out from the original data. As reference to gather gene and single nucleotide polymorphism (SNP) details from their HUGO Gene Nomenclature Committee (HGNC) Ids and rsids, a local copy of the Ensembl Human databases (version 75, databases core and variation, including SNPs coming from the 1000 Genome project) was employed; the annotation adopted for the whole analysis was GRCh37-p13, that includes the release 6 patches (Genome Reference Consortium: human assembly data - GRCh37.p13 - Genome Assembly (www.ncbi.nlm.nih.gov/assembly/GCF_000001405.25). We used ALIGATOR (Holmans et al. 2009; Houlston et al. 2010) program to evaluate how SNPs and single genes get summed to provide total contribution of candidate interactomes (Mechelli et al. 2013). The idea behind ALIGATOR’s strategy is to evaluate gene category significance by means of a bootstrapping approach: compare each interactome with the null hypothesis, that was built using random permutations of the data based on a non-parametric bootstrap analysis that uses the Gene Ontology database. Filtering criteria for the SNP selection included (i) a p-value significance taken at p-value cut-off (P-CUT) in the summary statistics; (ii) linkage disequilibrium (LD) filtering was applied. The parameters used to configure ALIGATOR are those reported in its reference paper (Holmans et al. 2009): P-CUT was taken at 0.05, the SNP-gene association parameter was maintained at its default value of 20kbp, and the LD filter was kept at r2<0.2. For all disease-associated interactomes, SNPs were extracted after the ALIGATOR filtering and bootstrapping analysis steps. The idea behind this extraction was to pinpoint which SNPs were considered relevant by the ALIGATOR algorithm. Genes (MS-AIG) were mapped to SNPs by the ALIGATOR internal mapping step, for bp distance, p-value, and LD, using the default settings optimized in Holmans paper (Holmans et al. 2009).

##### Protein binding enrichment

Genomic regions in Figure 1D were identified using Variant Effect Predictor version 98. After the ALIGATOR analysis, we used RegulomeDB database (Boyle et al. 2012), that is fed with live data from ENCODE and Roadmap Epigenomics, to look for proteins binding in regions containing “post-match” SNPs that displayed nominally significant associations with MS. When SNPs are bound by two or more proteins a Fisher’s exact test (2×2 table), controlling q-values for multiple-testing (FDR), Benjamini-Hochberg, was computed to evaluate whether this co-occurrence is statistically relevant using StatsModels v0.10 in Python v3.6.0.

##### Frequency of interactome genes in MS transcriptomes

The human PBMC microarray datasets analyzed in this study were published in Srinivasan papers (Srinivasan et al. 2017a; Srinivasan et al. 2017b) and deposited at EBI Array express database. The dataset was generated by Illumina Human Ref-8 v2 microarrays and contained PBMC transcriptomes of persons with MS (46 with clinically isolated syndromes, 52 with relapsing-remitting, 23 with primary-progressive, 21 with secondary-progressive disease) and 40 healthy controls.

The human brain microarray datasets were published (Magliozzi et al. 2019) and deposited at Gene Expression Omnibus. Briefly, the datasets were obtained by the Illumina whole genome HumanRef8 v2 BeadChip arrays, analyzing RNA extracted from chronic-active subpial GML (type III) and one nearby area of NAGM from the same tissue block in 10 cases of SPMS F+, 10 cases of SPMS F-, as well as in 10 control cases of non-neurological disease. The post-mortem tissues have been previously characterized for neuronal and glial alterations in cortical pathology, and for the presence of lymphoid-like immune cell aggregates (Magliozzi et al. 2010).

The enrichment of MS-AIG in DEG and in a random selection of transcript from the databases, was measured by chi-square test with Yates’s correction using Graph Pad.

##### Enrichment of EBV interactors in MS-AIG and HSP-protein modules

HSP-protein modules were identified by Bis-Brewer (Bis-Brewer et al. 2019) through a Protein-Protein interaction criteria using Human Integrated Protein-Protein Interaction rEference (HIPPIE) database and the DIseAse Module Detection (DIAMOnD). We performed an overlap between MS-AIG (n=741) and the Hereditary Spastic Paraplegia dominant-protein modules (n=295). To verify the apparent enrichment of EBV interactors in HSP modules, we randomly extracted from MS-AIG, 100.000 samples of the same size of shared genes and counted the number of EBV interactors included in each random sample. We considered the shared genes as enriched for EBV interactors, if the p value of the observed number under the null distribution was <0.05. The analysis was performed using R version 1.1.456.

##### Network and pathway analysis

MetaCore© (version 6.29 build 68613; GeneGO, Thomson Reuters, New York, N.Y) was employed to classify genes, carrying SNPs obtained from ALIGATOR analysis, in “canonical pathways” and evaluate their possible enrichment in specific categories. Canonical pathway maps represent a set of signaling and metabolic maps covering human biological processes in a comprehensive way. All maps are created by Thomson Reuters scientists by a high-quality manual curation process based on published peer-reviewed literature and validation experiments (Bolser et al. 2012).

##### Pi-index based analysis for MS-AIG

Looking at the therapeutic potential of MS-AIG, we measured their frequency in the top 150 positions of a recent published repository (Pi-Ranking) based on GWAS variants for several immunopathological traits, including MS (Fang et al. 2019). We verified whether the resulting gene lists was statistically enriched in each candidate interactome using chi-square test (Graph Pad).

##### Immunohistochemistry on post-mortem MS brain samples

Snap frozen tissue blocks from 10 post-mortem brains of secondary progressive MS cases (Figure 4, Table S2) and of 3 age-matched post-mortem controls (2 female/1 male) without neurological conditions have been analyzed. Brains were obtained at autopsy from the UK MS Society Tissue Bank at Imperial College, under ethical approval by the National Research Ethics Committee (08/MRE09/31). Six (*in Figure 4A) out of the 10 examined MS cases were also included in the previous study of Illumina gene expression analysis (Magliozzi et al. 2019) and in the ddPCR described below. Air dried, 10μm thick cryosections were rehydrated with PBS and immunostained with myelin oligodendrocyte glycoprotein (MOG) or major histocompatibility complex (MHC) class-II antibodies for neuropathological assessment of lesion presence and activity following previously published procedures (Magliozzi et al. 2010). Serial frozen sections, cut from the same blocks, were stained using a specific anti-CD40 rabbit polyclonal antibody (Santa Cruz). Double immunohistochemistry staining was performed combining antibodies specific for CD40 and markers MHC-II for microglia/macrophages. Antibody binding was visualised using peroxidase or alkaline phosphatase systems (Vector Labs, Peterborough, UK). For MOG immunostaining treatment with cold methanol was performed, whereas for CD40 and MHC-II cold ethanol fixation was used. Images were acquired using an Axiophot microscope (Carl Zeiss, Jena), equipped with a digital camera (Axiocam MRC) and analyzed using Axiovision 6 AC software.

##### Digital droplet PCR assays

RNA sample obtained from subpial cortical lesions from 6 post-mortem MS cases and from NAGM of 3 healthy controls matched and previously examined (Magliozzi et al. 2019), was further analysed to validate the CD40 gene expression. Briefly, five-hundred ng of total RNA was reverse-transcribed using Omniscript Reverse Transcription kit (Qiagen) following the manufacturer’s instructions. Each sample was analyzed by a multiplex ddPCR approach according to the manufacturer’s instructions, using predesigned gene expression assays for CD40 (Hs01002915_g1, FAM-labelled, Lifetechnologies) and GAPDH housekeeping gene (dHsaCPE5031597, HEX-labelled, Bio-rad).

Relative quantity of CD40 versus GAPDH gene expression was evaluated using QuantaSoft software, version 1.7 (Bio-Rad, Hercules, CA). For a given sample copies/microliter were calculated by averaging over the duplicate wells.

## QUANTIFICATION AND STATISTICAL ANALYSIS

The quantitation of protein levels (Figure 3) was performed by FlowJo v10.5 and expressed as median fluorescence intensity. Unpaired two-tailed T-test was used for the comparisons between groups showed in Figure 3B-E, in Figure 4A and in Figure 5B. The correlations between protein levels (Figure 3 F and G) were performed by a simple linear regression analysis. The enrichment of MS-AIG in DEG (Table 1 and 2) as well as the enrichment of MS-AIG in Pi-ranking (Figure 5) were measured by chi-square test with Yates’s correction. All the above analyses were performed using Graph Pad Prism 8.

Fisher’s exact test (2×2 table), controlling q-values for multiple-testing [false discovery rate (FDR), Benjamini-Hochberg], was used to evaluate the statistical relevance of protein co-occurrence by using StatsModels v0.10 in Python v3.6.0.

The mixed-effect multiple linear regressions analyses showed in Figure 3H and I were performed using R program, Version 1.1.456 (function “lmer” from “lme4” package). All the significances were considered when p< 0.05

## DATA AND CODE AVAILABILITY

Gene expression data obtained from peripheral blood leukocytes and brain are available respectively at EBI Array express database and Gene Expression Omnibus. GWAS data form MS are available at IMSGC; GWAS data form other diseases are available at WTCCC2.

---

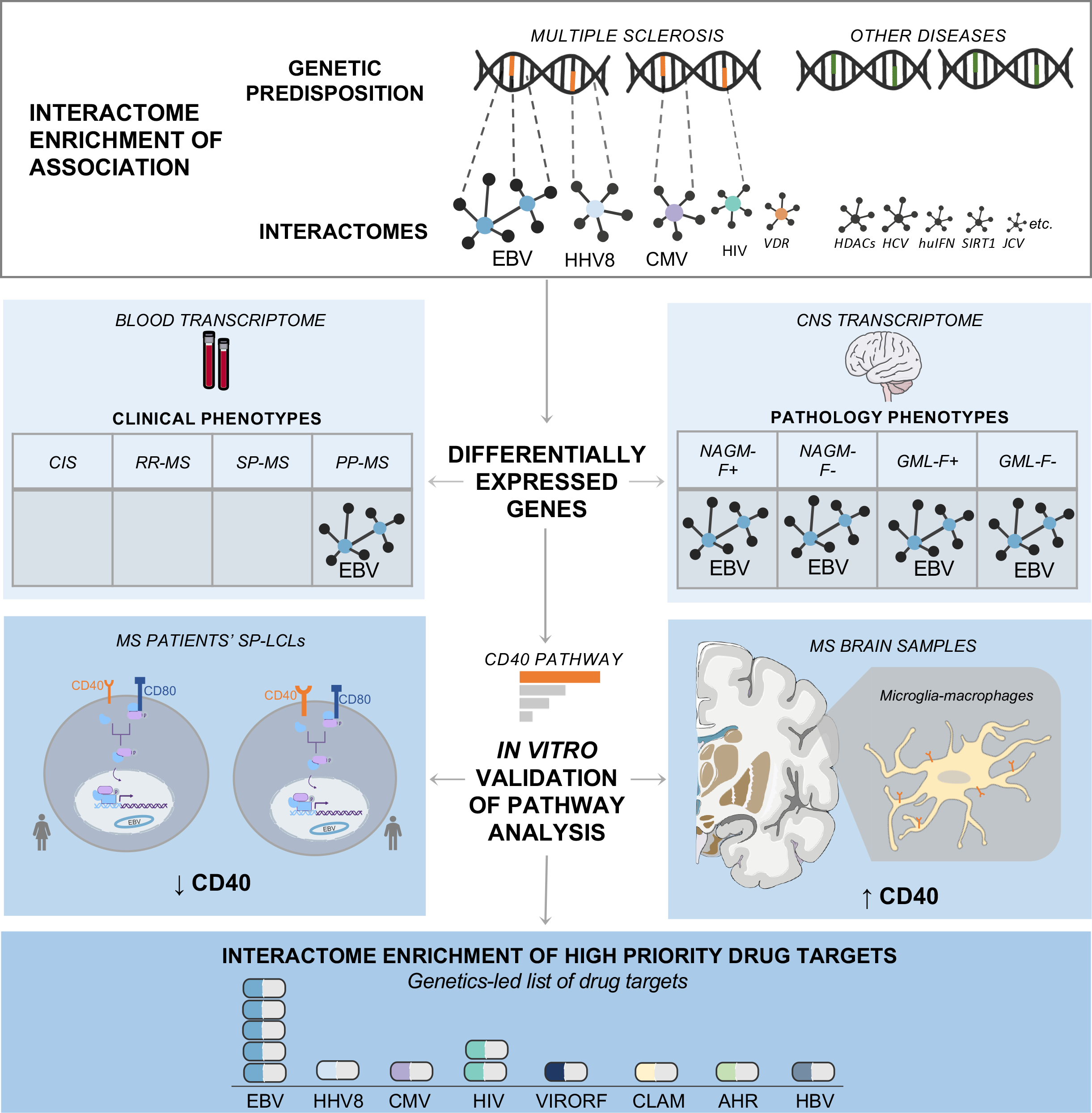

