## Supplemental informations_Mechelli et al. 2021 for "Enrichment analysis of GWAS data in autoimmunity delineates the multiple sclerosis-Epstein Barr virus association"

**GWAS**

**Immunochip**

**Metachip1**

chr1:pos1

chr1:pos2

chr1:pos3

chr1:pos1

chr1:pos2

chr1:pos4

chr1:pos1

chr1:pos2

chr1:pos3

chr1:pos4

chr1:pos1

chr1:pos2

chr1:pos3

chr1:pos4

**Metachip2**

GWAS SNPs

Immunochip SNPs

**Figure S1**: **Schematic representation of METACHIP datasets construction, related to Figure 1.**

Metachip 1 and 2 were built as the position-wise union of GWAS and Immunochip. Each dataset is represented by a line in which sample SNPs are indicated with different colors depending on their source dataset (black round for SNPs coming from GWAS and empty round for SNPs coming from Immunochip). When a given SNP existed in both GWAS and Immunochip, we gave preference to GWAS in Metachip1 and to Immunohip in Metachip2.

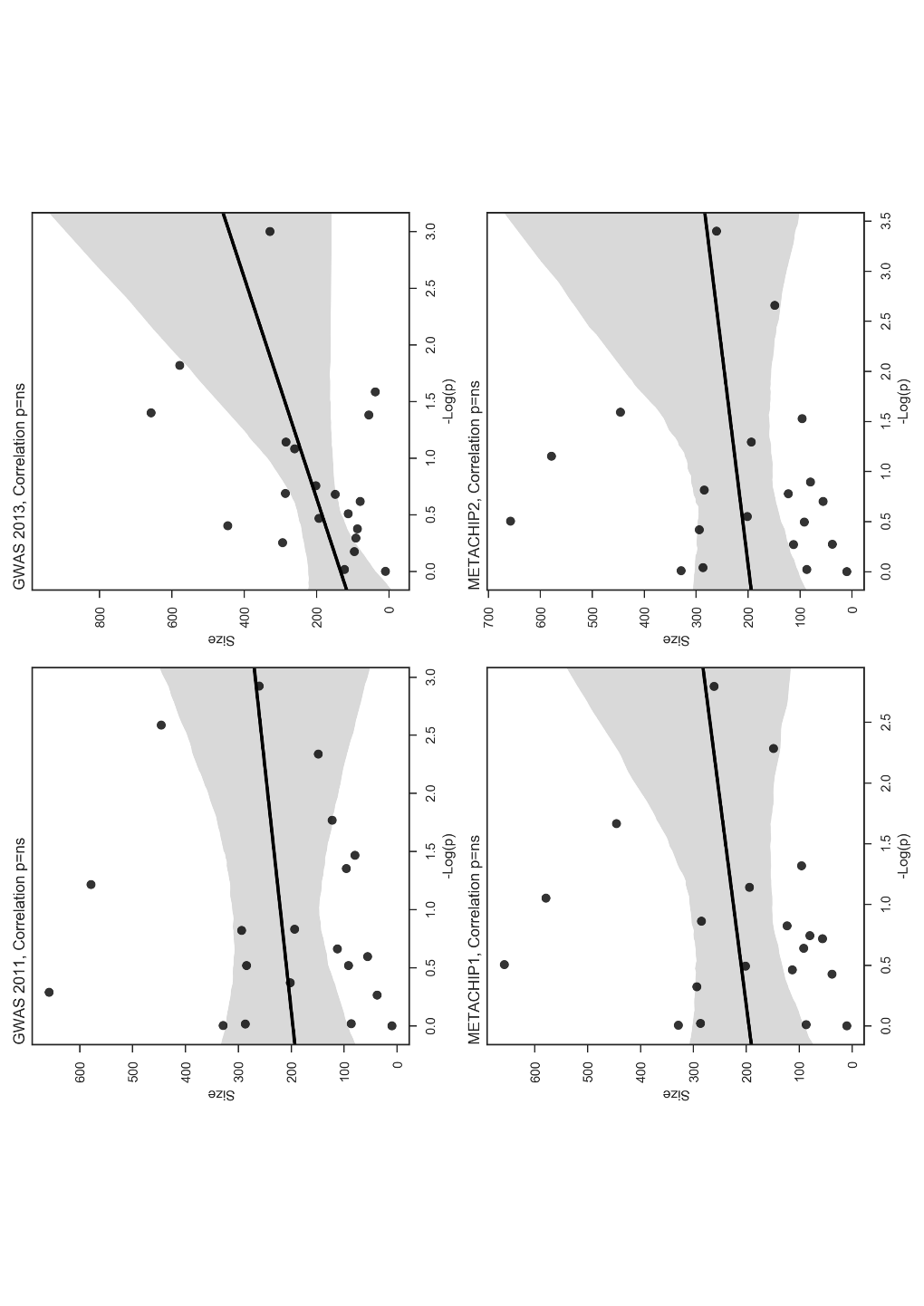

**Figure S2: Spearman correlation in MS GWAS, related to Figure 1.**

Linear correlations between the size of single interactomes and their cumulative p-value of association with MS calculated for each GWAS dataset (GWAS 2011, GWAS 2013, METACHIP 1 and METACHIP 2). The 95% confidence intervals of the Spearman correlation are greyed out in all the analyses.

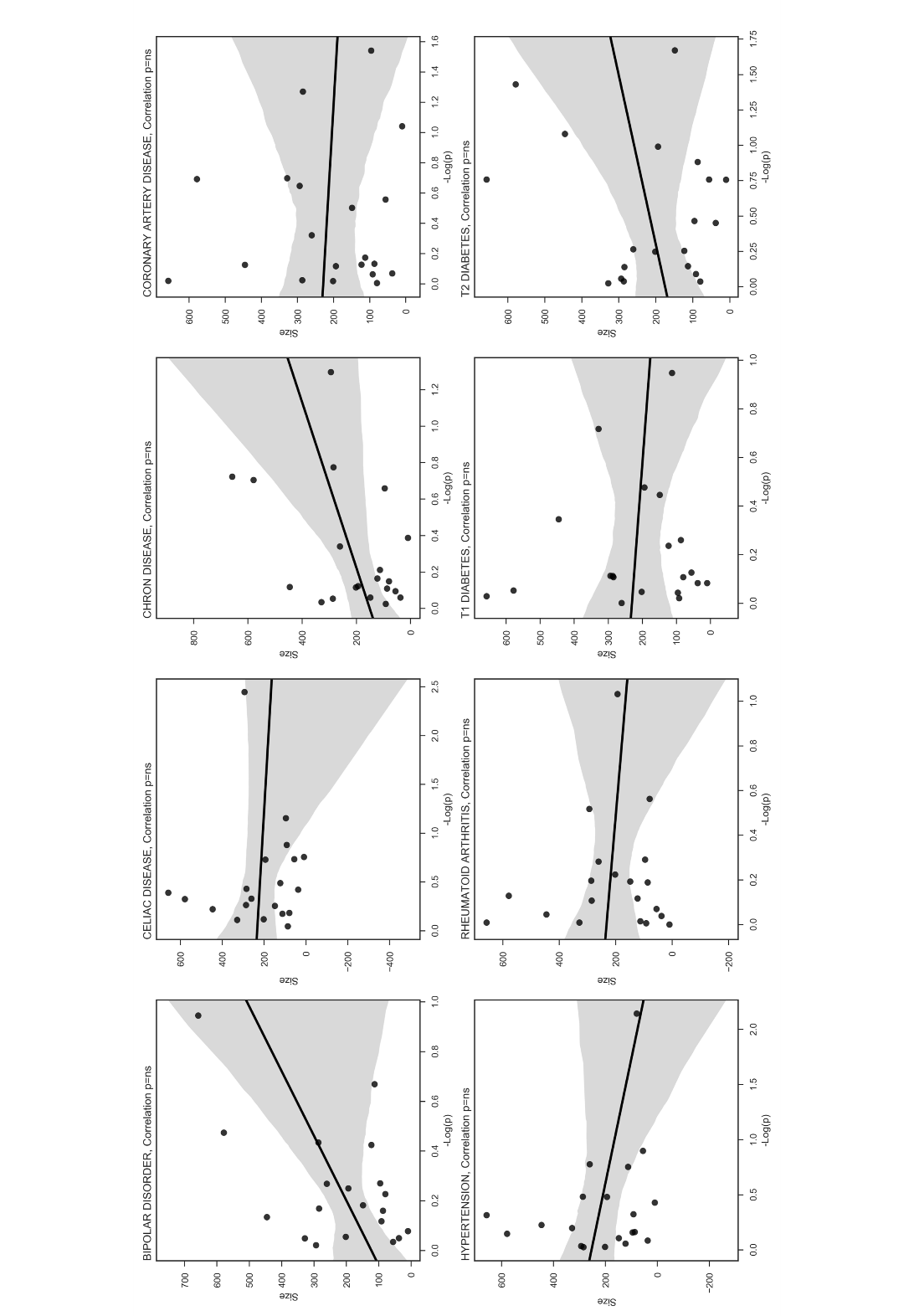

**Figure S3: Spearman correlation in non-MS GWAS, related to Figure 1.**

Linear correlations between the size of single interactomes and their cumulative p-value of association with complex diseases calculated for each GWAS dataset (Bipolar disorder, celiac disease, Chron disease, coronary artery disease, hypertension, rheumatoid arthritis, T1 diabetes, T2 diabetes). The 95% confidence intervals of the Spearman correlation are greyed out in all the analyses.

**Figure S4: Overlap between MS-AIG and genes of HSP-dominant expanded modules.**

(A) Overlap between MS-AIG and HSP-protein modules genes. Out of 71 shared genes, 34 resulted EBV interactors. (B) Null distribution of EBV interactors included in 100.000 random samples of the same size of the 71 shared genes, extracted from MS-AIG. The observed number of EBV interactors (N=34 out of 71 shared genes, red line) was larger than randomly expected (p value<0.00001).

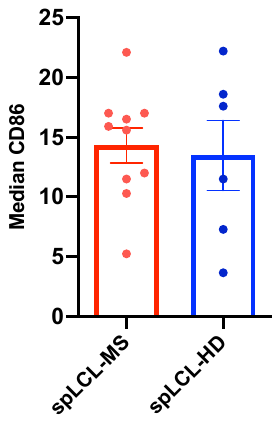

**Figure S5: The protein expression of CD86 on spLCL is comparable between MS (spLC L-MS, n=10) and controls (spLCL-HD, n=6; A), related to Figure 3.** The protein levels are represented as median fluorescence intensity. The data analysis was performed using Unpaired t-test.

ns= not statistically significant.

A	ns

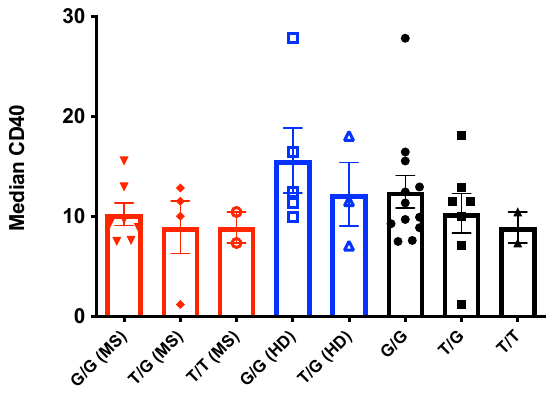

**All samples**

B	ns

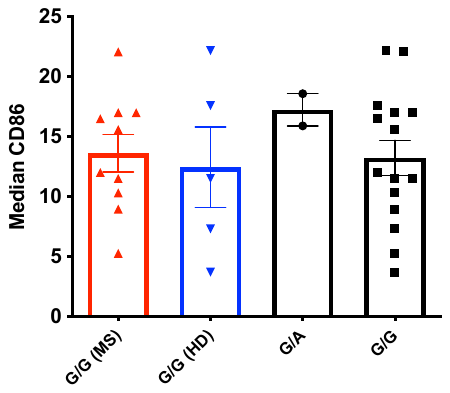

**All samples**

**Figure S6: Genotype independent CD40 and CD86 protein expression on spLCL, obtained from MS patients and controls (HD), related to Figure 3**. (A) The rs4810485*T is the CD40 multiple sclerosis risk allele. The protein expression levels (represented as median fluorescence intensity) are independent from the genotype when analyzed overall (GG, n=12; TG, n=7; TT, n=2) and when MS (GG, n=7; TG, n= 4; TT, n=2) and HD (GG, n=5; TG, n=3; TT =not detected) were analyzed separately.

(B) The rs9282641*G is the CD86 multiple sclerosis risk allele. The protein expression levels are independent from the genotype when analyzed overall (GG, n= 15; GA, n=2) and when MS (GG, n=10) and HD (GG, n= 5; GA, n=2) were analyzed separately.

The data analysis was performed using 1-way ANOVA, Bonferroni post-test. ns= not statistically significant

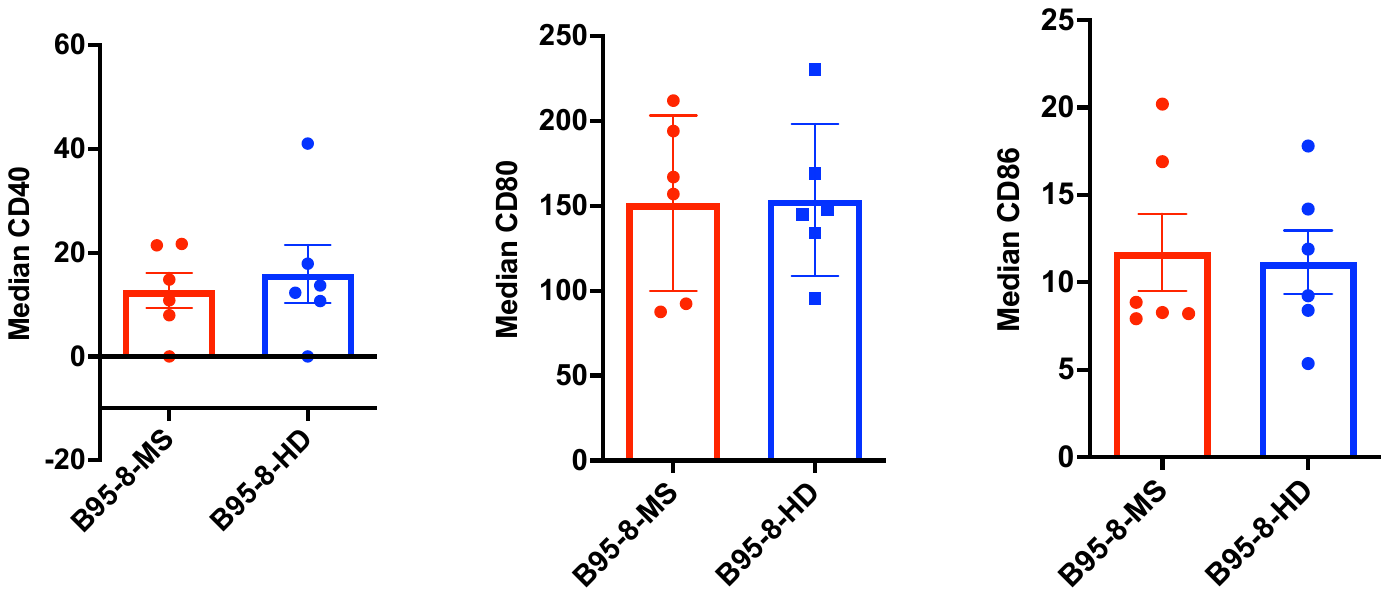

ns

ns

ns

**Figure S7: Protein expression of CD40, CD80 and CD86 on B95-8LCL surface, related to Figure 3.**

The protein levels are comparable between MS (n=6) and HD (n=6)-derived B95-8LCLs.

On *Y-axis* are represented the median fluorescence intensity of each protein. The data analysis was performed using Unpaired t-test.

ns = not statistically significant.

**Figure S8: Overlap between MS-AIG and the MS disease module derived from multi-omics analysis by Badam *et al,* 2021.**

(A) Overlap between all 741 MS-AIG and the 220 genes forming the MS module defined by Badam et al, 2021 through comparative multi-omics analysis.

(B) Overlap between the MS Multi-Omic Module and the MS-AIG differentially expressed in the brain, blood and both.

The overlap is considered significant if p <0,05 (Fisher’s exact test, calculated through *geneOverlap* package in R software). ns= not significant.

**Table S1: List of 741 MS-AIG, related to Figure 1**

| **MS-AIG BY INTERACTOME OF ORIGIN** | |
| --- | --- |
| **CMV** | *ACTL6A, CD209, CDC23, CDH1, CREB1, CSNK2A1, CSNK2B, DAXX, DDA1, DDB1, EGFR, EGR1, EP400, FKBP10, HDAC3, HNRNPA1, HNRNPH3, JUN, JUNB, KAT2B, KAT5, KPNA1, LILRB1, MICA, NFE2, NFE2L2, PDIA2, PDIA4, PML, PSMB4, PSMB6, PSMD2, PSMD3, RAB11FIP4, RAB1A, RUVBL2, SART3, SPI1, STAT1, STAT2, STAT3, TAP1, TAP2, TAPBP, TBP, TEAD1, TMEM43, TP53, TRAF6, TRRAP, TSC2, UBE2I, UBR5, USP7, VPRBP, ZMYND11* |
| **EBV** | *CSNK2A1, HNRNPA1, KAT5, KPNA1, PML, PSMB6, PSMD2, RUVBL2, SPI1, TAP1, TAP2, TRAF6, UBR5, USP7, ZMYND11, ACTB, AKAP8L, ALDH5A1, ATF7IP, C1QBP, CAD, CAMK2D, CAPZB, CCNA1, CDKN2A, CHEK2, CLTC, CR2, CTBP1, CUL3, CUL4A, CUL5, DARS, DDX17, DHX9, DNAJB6, EBNA1BP2, EIF2AK2, EIF3B, EP300, FLNC, FOXO1, GBAS, GRPEL1, H1FX, HDAC4, HIST1H1D, HIST1H2BG, HLA-DPA1, HLA-DPB1, HLA-DQA1, HLA-DQA2, HLA-DQB1, HLA-DRA, HLA-DRB1, HLA-DRB5, HMGB2, HNRNPAB, HNRNPK, HNRNPM, HNRNPR, HNRNPU, HSPA1A, HSPA4, HSPA8, HSPB1, IARS, IDH3A, IDH3B, ILF3, ING4, IQGAP2, IRF7, JAK3, KHDRBS1, KPNA2, KPNB1, LDHB, LRPPRC, LYN, MLF2, MRPS18B, MYC, NAP1L4, NEDD4L, NIPSNAP1, NUP93, P4HA1, P4HA2, PCBP1, PCBP2, PCCA, PCNA, PHB2, PRMT1, PRMT5, PRPF8, PSMC2, PTBP1, PYCR2, QARS, RAD18, RAD50, RBM15B, RCN1, RFC3, RIPK1, RNF4, RPA1, RPL11, RPL13, RPL18, RPL19, RPL22, RPL24, RPL27A, RPL34, RPL7A, RPL9, RPS13, RPS18, RPS2, RPS25, RPS6, RPS8, RPS9, SEC13, SEC16A, SERBP1, SF3B2, SNRNP200, SNRPA, SNRPD2, SPEN, SRC, STUB1, SYK, TERF2, TNFAIP3, TRAF1, TRAF2, TRAF3, TUBB, TYK2, VDR, XPOT, XRCC5,* |
| **HHV8** | *ACAP2, ASPSCR1, C5orf17, CARM1, CDK2AP1, CDK6, CDKN1B, CHEK1, CIAO1, CREBBP, CUL4A, CUL5, DDX39B, DPPA2, EP300, ERI1, FBXW7, GFPT2, GSK3B, HCFC2, HIST1H2AI, HIST1H2AK, HIST1H2AL, HIST1H2AM, HIST1H2BC, HIST1H2BE, HIST1H2BF, HIST1H2BG, HIST1H2BI, HIST1H4A, HMGB1, HNRNPK, HNRNPUL1, IKBKB, IRF1, IRF6, IRF7, JUN, KAT5, MAP3K7, MED16, MED24, NCOA2, NFKBIA, NRF1, OLIG3, PARP1, PCBP2, PCNA, POLB, POU2F1, PPIE, PPP2R1A, PPP5C, PRKDC, PRMT1, PRMT2, PRMT3, PRMT5, PRMT6, PRMT7, PRMT8, RAD51C, RBM19, REEP2, RELL2, RGS14, RIPK1, RPA1, RPA2, RRM1, SERBP1, SMARCA4, SMARCC2, SND1, STAT3, STAT5A, SUMO3, TCEB1, TCEB2, TERF1, THOC5, TLE2, TP53, TRAF2, UBE2D1, UBE2I, USP7, VHL, WWP1, ZBTB38* |
| **HIV** | *ACBD5, ACSL3, AFF1, AIMP2, AKAP8L, ALMS1, ANAPC1, APP, ATP2A2, ATP2A3, ATP2B1, ATP5H, ATP5J, ATXN10, CALU, CANX, CAPN1, CARM1, CASC3, CDKN2A, CERS2, CHAF1B, CHCHD1, CHI3L2, CIAO1, CLN6, CNTNAP1, COMT, COPS3, COPS4, COPS6, COX4I1, COX5A, COX6B1, CUL5, DAGLB, DCAF8, DDA1, DDX6, DIAPH1, DNAJB11, DNAJC3, DNAJC7, DYNC1H1, DYNC1I2, EEF1E1, EFEMP1, EIF3A, EIF3B, EIF3D, EIF3E, EIF3K, ENDOD1, ERGIC1, ERGIC2, FAM203A, FAM98A, FBN2, FBXO3, FBXW11, FLCN, FLOT1, FOXC1, G3BP1, GEMIN2, GIMAP5, GLT25D1, GRB2, HADHB, HAT1, HDAC3, HIST1H3A, HIST1H3B, HIST1H3C, HIST1H3D, HIST1H3E, HIST1H3F, HIST1H3G, HIST1H3H, HIST1H3I, HIST1H3J, HLA-B, HNRNPH3, HSD17B12, HSP90B1, HSPH1, IDH3A, IFT122, IK, ITGAL, ITGB2, KIAA1429, KIAA2013, KIF21B, KIR3DL1, KIR3DL2, LARP7, LDLR, LIMD1, LMAN2, LRPPRC, LRRC47, LSM12, MARS, MCL1, MCM7, MEPCE, MLLT1, MTHFD2, NAP1L4, NEDD4L, NFU1, NMT1, NUDC, OLA1, OS9, OSBPL6, P4HA2, PDIA4, PDIA5, PLOD1, PLOD2, POLE, PPP1R12A, PRKDC, PRMT1, PSMA1, PSMB6, PUM2, RABGAP1, RANBP2, RCN1, RNF126, RNF7, RNH1, RTN4, SDCBP, SDHA, SKP1, SLC27A4, SMU1, SPCS3, SUMF2, SUN1, SURF4, TBL1XR1, TCEB1, TIMM13, TMEM160, TOMM40, TRPS1, UBE2O, UNC45A, VPRBP, XPO1, XPO4, YTHDF3, YWHAB, YWHAZ* |
| **HBV** | *BUB3, CAK, CD44, CD81, CFLAR, CREB1, DBI, DDB2, DNMT3A, EGR1, EP300, ERCC3, FETUB, GTF2H3, GTF2H4, GTF3A, HIF1A, HNF4A, HSP90AB1, HSP90B1, HSPA1A, HSPB1, IL6, JAG1, JAK1, L3MBTL3, MAP3K1, MAPK1, MAPK8, MAPK9, MAVS, MPDU1, NFKB1, NFKBIA, NFKBID, NFKBIE, POU2F1, PPP2CA, PRMT1, RARA, SKP2, SLC25A23, SREBF1, SRPK1, SRPK2, STAT1, TM4SF4, TPSB2* |
| **AHR** | *ARNTL, DAP3, DDB1, KIAA1683, MAF, NCOA1, NCOA2, NRIP1, RELA, SMARCA4, SRC, STUB1, TBP* |
| **POLYOMAVIRUS** | *BTRC, CDK2, CSNK2A1, HIST1H1A, HIST1H1T, IRS1, KIN, KPNB1, LTA, PARK7, PARP1, PPP2CA, PPP2R1B, RPA1, STUB1, TBP, TEAD1, TOP1* |
| **SIRT7** | *ACTR2, AHNAK, ANKHD1, ANKRD52, ANXA6, APP, ARID1A, ATAD2B, ATAD3A, ATM, ATXN2L, BEND3, CCT5, CHD4, CHERP, CKAP5, COPA, COPB1, CSNK2A1, DDB1, DHX16, DLD, DNM2, DNMT1, DST, EBNA1BP2, EDC4, EIF4G3, EPB41L3, FUS, FXR2, GANAB, GEMIN4, GFPT1, GTF2I, GTPBP4, H1FX, HDLBP, HIST1H1A, HIST1H2BL, HIST1H3A, HP1BP3, HYOU1, IMPDH2, INTS1, IQGAP1, ITPR3, KDM1A, KIAA0020, KIF1A, KPNB1, KRI1, LARP4B, LMNA, LTA4H, LTF, MCM6, MDC1, MLL, MPHOSPH10, MTOR, MYBBP1A, NCAPD2, NOC2L, NOL9, NOP2, NPEPPS, NR0B2, NSD1, NSUN2, NUP210, NUP98, OASL, PABPC4, PCNT, PFKP, PHRF1, PLEC, PLOD1, POLR2A, PRMT5, PRRC2A, PUF60, RAD50, RANGAP1, RBM6, RCC2, RCL1, RNH1, RPL3, RPL5, RPL6, RPLP2, RPS26, RRP8, SCAF11, SDHA, SEC13, SEC16A, SEC24C, SERBP1, SKIV2L, SMARCA2, SMARCA4, SMARCC2, SND1, SPEN, SRP68, SRRM1, SRRM2, SYMPK, TJP1, TOP1, TSR1, UQCRC1, USP34, USP47, UTP20, WDR12, WDR46, WDR6, WHSC1, XPOT, ZC3HAV1, ZCCHC3* |
| **VIRORF** | *ACLY, ACP1, ANP32B, ATAD3A, ATXN2L, BASP1, C10orf32, C10orf90, CALM1, CASP8, CCT5, CDC37, COPB1, CSDA, CSNK2A1, CSNK2B, CTBP1, CUL4A, DDB1, DDX39B, DDX6, DNAJB11, EDC4, EEF1A2, ELANE, ENO1, FUS, GAPDH, GNB1, GNB4, GRPEL1, GTF2I, H1FX, HIST1H1A, HIST1H1T, HIST1H2AB, HIST1H2AC, HIST1H2AD, HIST1H2AJ, HIST1H2BA, HIST1H2BC, HIST1H2BD, HIST1H2BL, HIST1H2BM, HIST1H2BN, HIST1H2BO, HIST1H3A, HNRNPAB, HNRNPK, HNRNPU, HSPA1L, HSPA6, HYOU1, IFIH1, IKBKB, ILF3, KDM1A, KEAP1, KHNYN, KPNB1, LDHB, LEMD2, LENG8, LTF, MAD1L1, MAP4K4, MIB2, MYBBP1A, MYL6, MYL6B, NAP1L4, NUP98, NYNRIN, PABPC4, PCBP3, PFKP, PIK3R2, PIK3R3, POLD1, POLR2A, POM121C, PPP2CA, PRTN3, PTBP1, ROCK2, RPL18, RPL22, RPL27A, RPL3, RPL32, RPL36, RPL5, RPL6, RPS18, RPS20, RPS21, RPS26, RPS9, SEC24C, SEC31A, SERBP1, SF3B3, SLC25A3, SMARCA4, SMARCC2, SMARCE1, SMYD3, SOLH, SPOP, STAT1, STAT2, STUB1, TCEB2, TRAF6, TRIM21, TSSC1, TUBB, TUBB2A, TUBB2B, USP19, USP24, USP54, VAPA, VPS37B, VPS52, VPS53, WDR6, WRNIP1, YWHAQ, ZMYND11* |
| **CLAM** | *ALG1, AP3D1, ARRB2, ATP5I, ATP6V0A1, CAND2, CAPNS1, CDKAL1, COPS7A, CSNK2A1, CSNK2B, DAP3, DAXX, DCTN2, DCTN4, DCTN5, DIAPH1, DNAJC11, DPP9, EPB41L3, GNB4, GTF2I, HEATR3, HLA-A, HYOU1, IDE, KDSR, KIR2DL4, LMAN2, MAP4K4, MAPRE1, MCM6, MRPS18B, MRPS22, MTX1, NANS, NCAPH2, NOTCH1, NOTCH2, PIP4K2C, PPP4C, RAB1A, RAB8B, RELA, RER1, RFT1, RNH1, RRBP1, SCAMP3, SCO2, SEC62, SIGMAR1, SIGMAR1, STX18, SYNGR1, TAOK2, TMED1, TMEM106B, TTC35, UBR2, VANGL2, VAPA, VMP1, WDR6, YWHAH, ZZEF1* |
| **AIRE** | *ASAH1, CHD6, CREBBP, CREG1, CTSH, DAXX, DDX17, EFTUD2, FAT1, GDPD3, GM2A, HIST1H2AC, HIST1H2BO, HIST1H3A, HSP90AB1, HSPA1A, HSPA1L, HSPA4, KPNB1, MCM2, MSH2, MSH6, NUP93, PARP1, PRKDC, RANBP2, RAVER1, RUVBL2, SRSF2, TGM1, TMPRSS13, TUBA1C, TUBB, XPO1* |

**Table S2: Demographic, clinical and neuropathological data of the examined progressive MS cases. The presence (+) or absence of detected lesion is shown, n.a. indicated non-available lesions in the examined MS cases. Cases indicated with asterisks (*) were also included in the Illumina and digital droplet-PCR (ddPCR) gene expression analysis displayed in Figure 4.**

|  |  |  | |  | **GM** | | | | **WM** | | | |
| --- | --- | --- | --- | --- | --- | --- | --- | --- | --- | --- | --- | --- |
| **Cases MS** | **Sex** | | **Age a death**  **(years)** | **Disease duration**  **(years)** | **NAGM** | **Active lesion** | **Chronic active lesion** | **Inactive lesion** | **NAWM** | **Active lesion** | **Chronic active lesion** | **Inactive lesion** |
| **MS74*** | **Female** | **64** | | **36** | **+** | **n.a.** | **-** | **n.a.** | **+** | **n.a.** | **-** | **-** |
| **MS141** | **Male** | **66** | | **37** | **-** | **+** | **n.a.** | **n.a.** | **-** | **+** | **-** | **n.a.** |
| **MS163*** | **Female** | **45** | | **6** | **-** | **n.a.** | **-** | **-** | **+** | **n.a.** | **-** | **-** |
| **MS200*** | **Female** | **44** | | **19** | **-** | **+** | **+** | **-** | **-** | **+** | **n.a.** | **-** |
| **MS311** | **Female** | **45** | | **16** | **-** | **n.a.** | **n.a.** | **n.a.** | **+** | **n.a.** | **n.a.** | **n.a.** |
| **MS79*** | **Female** | **49** | | **21** | **+** | **n.a.** | **-** | **n.a.** | **+** | **+** | **-** | **n.a.** |
| **MS121*** | **Female** | **49** | | **14** | **-** | **n.a.** | **-** | **n.a.** | **-** | **+** | **-** | **-** |
| **MS136*** | **Male** | **40** | | **12** | **-** | **+** | **n.a.** | **-** | **+** | **n.a.** | **-** | **n.a.** |
| **MS286** | **Male** | **45** | | **16** | **+** | **n.a.** | **-** | **-** | **-** | **+** | **+** | **-** |
| **MS330** | **Female** | **59** | | **39** | **-** | **+** | **+** | **-** | **-** | **n.a.** | **n.a.** | **n.a.** |

**Table S3: MS-AIG targeted by drugs (with their trial phase and indication) resulting from Fang et al. 2019, related to Figure 5.**

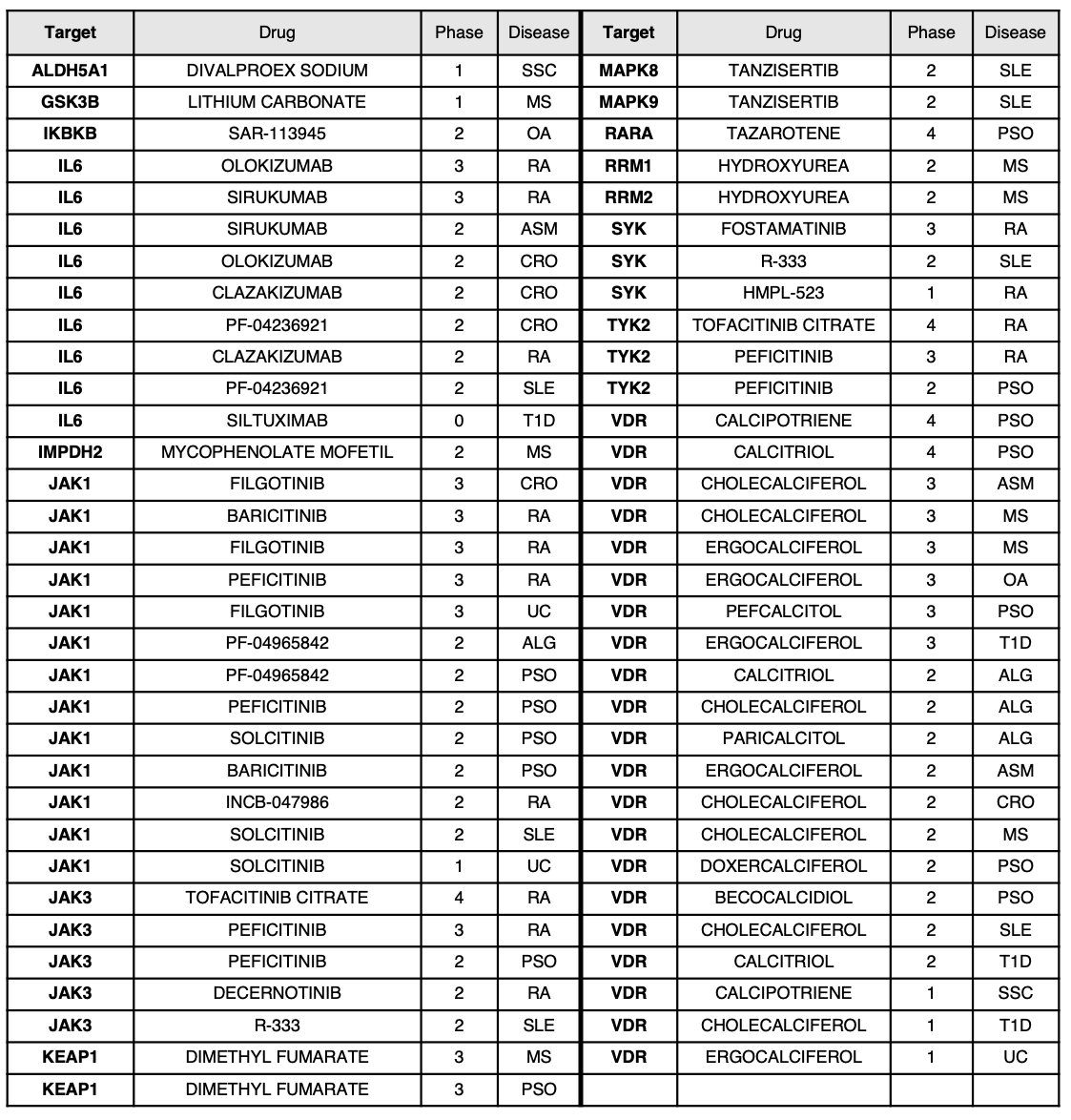

ALG = Allergy, ASM = Asthma, CRO = Crohn’s disease, MS = Multiple Sclerosis, OA = Osteoartrhitis, PSO = Psoriasis, RA = Rheumatoid Arthritis, SLE = Systemic Lupus Erythematosus, T1D = Type 1 Diabetes, UC = Ulcerative Colitis
